# CRISPR-Cas9 mediated nuclear transport and genomic integration of nanostructured genes in human primary cells

**DOI:** 10.1101/2021.11.08.467750

**Authors:** Enrique Lin-Shiao, Wolfgang G. Pfeifer, Brian R. Shy, Mohammad Saffari Doost, Evelyn Chen, Vivasvan S. Vykunta, Jennifer R. Hamilton, Elizabeth C. Stahl, Diana M. Lopez, Cindy R. Sandoval Espinoza, Alexander E. Dejanov, Rachel J. Lew, Michael G. Poirer, Alexander Marson, Carlos E. Castro, Jennifer A. Doudna

**Author notes:** Equal contributions (Enrique Lin-Shiao, Wolfgang G. Pfeifer).

## Abstract

DNA nanostructures are a promising tool for delivery of a variety of molecular payloads to cells. DNA origami structures, where 1000’s of bases are folded into a compact nanostructure, present an attractive approach to package genes; however, effective delivery of genetic material into cell nuclei has remained a critical challenge. Here we describe the use of DNA nanostructures encoding an intact human gene and a fluorescent-protein encoding gene as compact templates for gene integration by CRISPR-mediated homology-directed repair (HDR). Our design includes CRISPR-Cas9 ribonucleoprotein (RNP) binding sites on the DNA nanostructures to increase shuttling of structures into the nucleus. We demonstrate efficient shuttling and genomic integration of DNA nanostructures using transfection and electroporation. These nanostructured templates display lower toxicity and higher insertion efficiency compared to unstructured double-stranded DNA (dsDNA) templates in human primary cells. Furthermore, our study validates virus-like particles (VLPs) as an efficient method of DNA nanostructure delivery, opening the possibility of delivering DNA nanostructures *in vivo* to specific cell types. Together these results provide new approaches to gene delivery with DNA nanostructures and establish their use as large HDR templates, exploiting both their design features and their ability to encode genetic information. This work also opens a door to translate other DNA nanodevice functions, such as measuring biophysical properties, into cell nuclei.

**Teaser Sentence:** CRISPR-Cas9 mediates nuclear transport and integration of nanostructured genes in human primary cells

## Introduction

Programmed self-assembly of DNA nanostructures (*1*–*5*) has applications in nanomanufacturing (*6*), biosensing (*7*) and biophysics (*8*–*10*). While delivery of DNA nanostructures to cells was one of the first proposed applications, progress has been hampered by challenges including uptake and stability of structures in cells (*11*). Previous work showed that DNA nanostructures remain stable in cell lysate for up to 24 hours and a single study has demonstrated cytosolic delivery with electrotransfection (*12*).

The scaffolded DNA origami approach (*3*), is particularly well suited to package sequences of several kilobases into a compact nanostructure. Because these DNA nanostructures are agnostic to the underlying DNA sequence, structures that exploit both their design features and their ability to encode genetic information can be engineered. This offers a promising route for nanostructure-mediated gene delivery.

While previous efforts have demonstrated the ability to effectively deliver small molecules (*13*, *14*), peptides and proteins (*15*, *16*), and small nucleic acids like siRNA (*17*, *18*) to cells, these studies require delivery to the cell surface or cytoplasm. Delivery of genomic information to the nucleus has remained a key challenge. Ideally, a nanostructure gene delivery system could target a gene to a specific genome site for integration.

CRISPR-Cas9 homology-directed repair (HDR) thus offers an attractive route since the gene of interest can be targeted through the inclusions of homologous sequences of DNA, which are straightforward to include on a DNA nanostructure. Furthermore, DNA nanostructures could offer a route for developing improved HDR templates for genome engineering. For example, long single-stranded DNA (ssDNA) donor templates can be folded to co-localize terminal sequences bearing homology to the intended genomic insertion site (homology arms). In addition, nanostructures can create compaction that could improve cellular delivery, increase half-life and circumvent toxicity of free DNA. Despite this promise for gene delivery with DNA nanostructures, several key advances are yet to be demonstrated including packaging of genes, effective delivery to the nucleus of live cells, integration of nanostructured genetic material into the genome, and targeting exogenous genes to a genome site of interest.

In this study, we tested strategies for nuclear delivery of DNA nanostructures encoding genes that can be used as HDR donor templates for precise, large genomic insertions using CRISPR-Cas9. Comparison of different methods for DNA introduction into cells showed electroporation to be an effective delivery strategy, providing the first evidence for nuclear localization of nanostructures. DNA nanostructures with short terminal sequences matching the sequence of the genomic integration site increased genomic insertion efficiency induced by Cas9 ribonucleoproteins (RNPs). This strategy was used in primary human T cells to replace a defective copy of *IL2RA*, a gene mutated in a familial immune dysregulation syndrome. We also tested Cas9 virus-like particles (Cas9-VLPs) for the co-delivery of DNA nanostructures, finding that relative to unstructured DNA templates, nanostructured DNA templates doubled the observed Cas9-induced genomic integration efficiency. These results demonstrate the utility of DNA nanostructures for some applications of genome editing and suggest that DNA template structure may assist both the delivery and use of DNA in other therapeutic and bioimaging applications. Furthermore, the ability to deliver DNA nanostructures to cell nuclei opens a door to translate other functions of DNA nanotechnology to the nucleus, such as force sensing (*19*), molecular detection (*20*), and biophysical measurement (*21*).

## Results

### Design of a DNA nanostructure encoding mNeonGreen for human genome integration

To test whether DNA nanostructures can enter the nucleus, and whether folding into a compact DNA architecture affects template utility for integration into the human genome, we designed a 2716-nucleotide ssDNAscaffold encoding mNeonGreen (*22*) (Fig. 1A, B, Fig. S17, 28). At each end, 100-nucleotide homology arms matched the sequences flanking the intended CRISPR-Cas9 cleavage site, such that the final integrated new sequence would be 2516 base pairs following successful HDR. In addition to mNeonGreen, the insertion segment includes a transcriptional promoter, a polyadenylation signal and a woodchuck hepatitis virus post-transcriptional regulatory element (WPRE). Successful integration of the template yields green fluorescent cells (mNeonGreen+), enabling detection of genome integration events.

**Figure 1:**
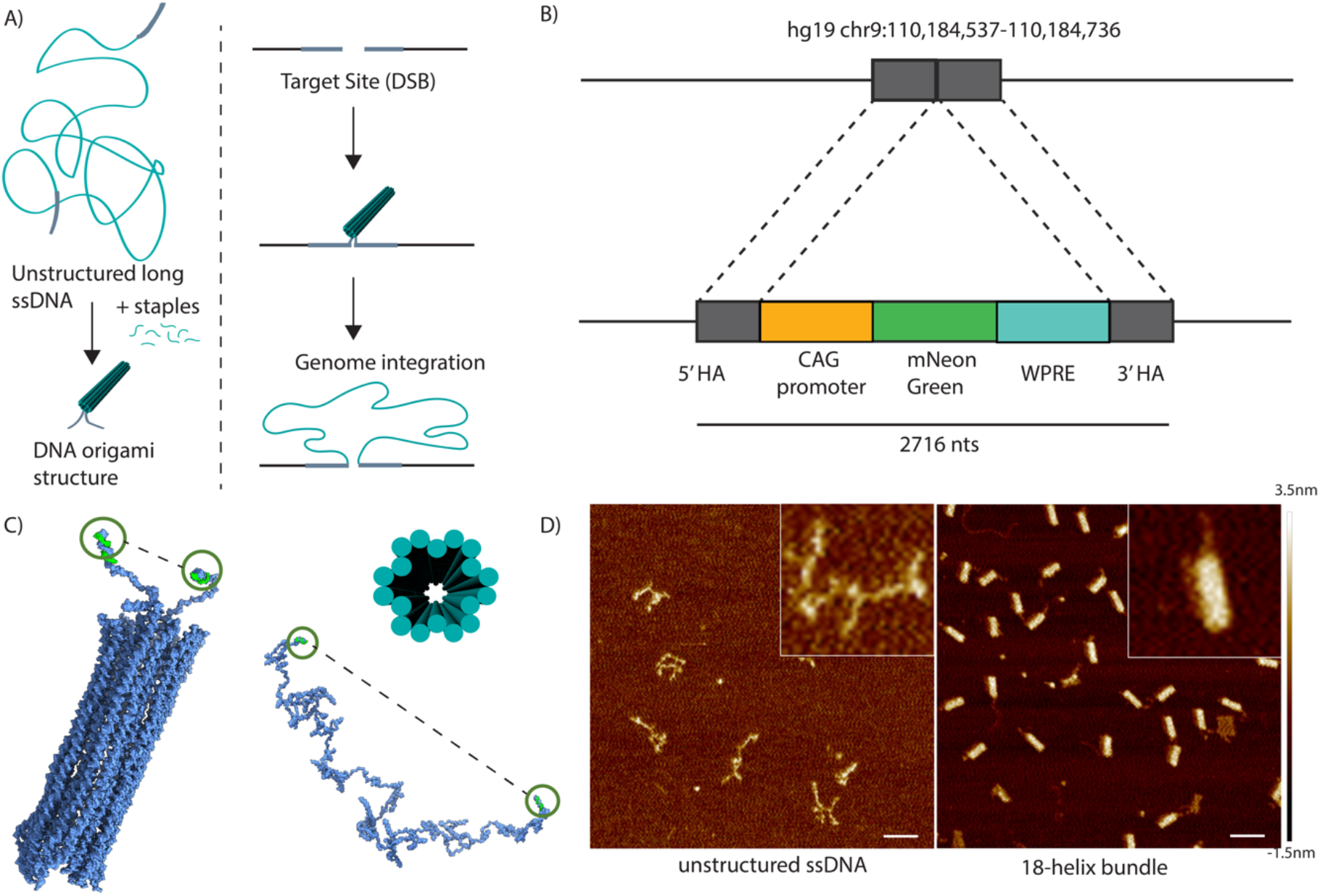
DNA nanostructure encoding mNeonGreen for human genome integration. A) Graphical strategy depiction showing folding of a long unstructured ssDNA into a DNA nanostructure for integration into the genome via CRISPR-Cas9 mediated HDR. B) Schematic of a 2716-base long template encoding mNeonGreen along with regulatory elements and two 100 bases homology arms for genome targeting atan intergenic site on human chromosome 9. C) Cylindrical model and oxDNA simulations of an 18 helix bundle DNA nanostructure show a decrease in end-to-end distance from 108.98 ± 11.22 nm (ssDNA) to 29.33 ± 9.9 nm (18 helix) D) AFM characterization of the unstructured ssDNA and the 18 helix DNA nanostructure. Scale bar: 100 nm.

Following established DNA nanostructure design rules, we created the mNeonGreen integration template as a hollow 18 helix bundle (herein called “18-helix nanostructure”) (*3*, *23*, *24*) (Fig. 1C). We performed molecular dynamics simulations (Movie S7, 8) implementing the coarse-grained model oxDNA (*25*, *26*) to guide the design process, assessing folded-structure energetics and comparing distances between DNA template termini. Consistent with expectations, the oxDNA simulations predicted that terminal homology sites are farther apart in the unstructured template (109 ± 11 nm) compared to the folded nanostructure (29 ± 10 nm) (Fig. S31-34, Table S3). Analysis by native gel electrophoresis showed that the 18-helix nanostructures migrated as a single species in each case, indicating correct structural formation (Fig. S35). Atomic force microscopy (AFM) revealed compacted, uniform DNA structures consistent with the designed properties of the 18-helix nanostructures (Fig. 1C, D). Conformations observed in AFM were consistent with simulations for both the unstructured template and folded nanostructures (Fig. S17-20, 28-30).

### Nuclear localization and genome integration of nanostructured DNA

After confirming the 18-helix nanostructures were folded, we next tested whether they can enter the nucleus and integrate into the human genome following genome cleavage by CRISPR-Cas9. Two different DNA delivery strategies were employed using human embryonic kidney 293T cells (HEK293T). First, we transfected 0.5 pmoles of 18-helix nanostructures together with plasmids encoding CRISPR-Cas9 and a single-guide RNA (sgRNA) targeting the aforementioned site on chromosome 9 using Lipofectamine (Fig. 1B; Fig. 2A). In parallel, we electroporated HEK293T cells with CRISPR-Cas9 RNPs together with 0.5 pmoles of 18-helix nanostructures. Both experiments included controls in which either 0.5 pmoles of unstructured ssDNA (herein called “unstructured”) or 0.5 pmoles of a simple DNA nanostructure where the ends of the ssDNA are connected together through base-pairing of several strands to fold the template into a closed loop (herein called “looped”) were used in place of 18-helix nanostructures to provide the HDR template during genome editing (Fig. 2A). After seven days, cells were harvested and analyzed by flow cytometry to assess the percentage of mNeonGreen-positive cells. Results showed that using both modes of delivery, DNA nanostructures can enter the nucleus and become integrated in the genome. However, 18-helix nanostructures delivered by transfection resulted in decreased HDR efficiency relative to unstructured DNA (<2% versus ~5%) (Fig. 2B, C). When using electroporation, HDR levels were similar for both 18-helix nanostructure templates and unstructured templates (4% versus 5.5%, respectively) (Fig. 2B, C). Interestingly, the closed looped nanostructure, in which homology arms are proximal but the template itself is unstructured, resulted in a slight increase in HDR efficiency in both transfected (~5%) and electroporated (7%) samples, although this result was not consistently reproducible (Fig. 2B, C). We used primers flanking the Cas9 cleavage site to confirm insertion of the mNeonGreen construct at the expected position (Fig. 2D). To determine whether electroporation affects DNA nanostructural integrity, we diluted 18-helix nanostructures in electroporation media and electroporated half of the sample. AFM revealed that DNA nanostructures maintained integrity after electroporation (Fig. 2E). These results suggest that DNA nanostructures can be delivered into the nucleus through electroporation and serve as templates for CRISPR-Cas9-mediated HDR.

**Figure 2:**
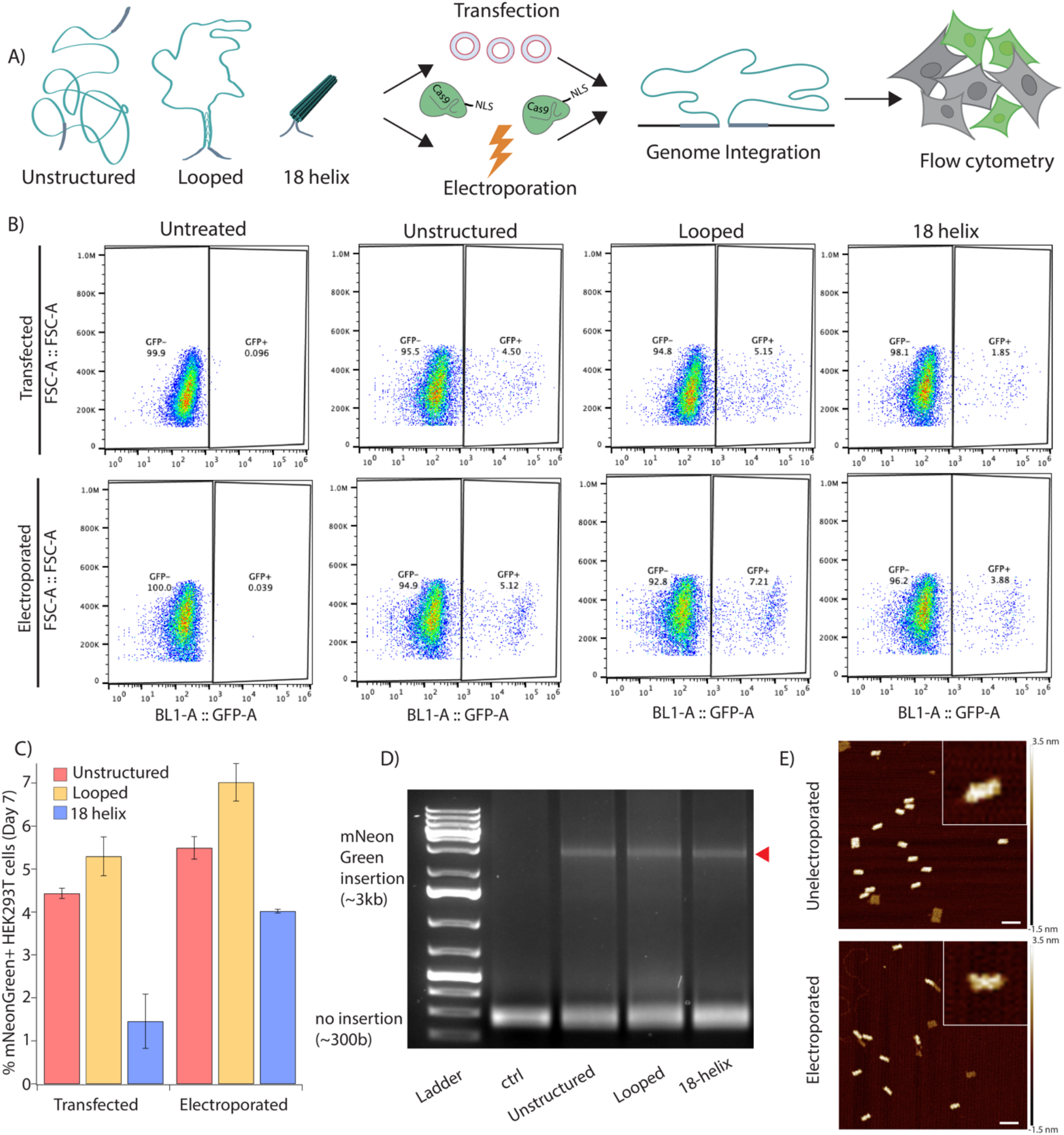
Nuclear localization and genome integration of nanostructured DNA. A) Schematic of experimental approach. B) i. Flow Cytometry data measuring mNeonGreen+ cells (GFP+) shows looped templates are more efficiently incorporated into the genome compared to unstructured and 18 helix nanostructures. ii. Flow cytometry of electroporated cells shows similar values across unstructured, looped and 18 helix nanostructures. C) Aggregated flow cytometry data shows looped templates perform best for both transfection and electroporation. Error bars represent SDs from 3 experiments. D) PCR using primers flanking the insertion site confirms mNeonGreen insertion at the target site (red triangle). E) AFM images of the 18 helix nanostructure before and after electroporation. Scale bar: 100 nm.

### Increased HDR efficiency upon CRISPR-Cas9 RNP localization at template DNA ends

These results demonstrate that electroporation is effective at delivering nanostructured DNA to the nucleus. We next investigated whether adding truncated Cas9 target sequences (shuttles) to DNA nanostructures could enhance the rate of nuclear localization and subsequent genome integration (*27*, *28*) (Fig. 3A). Our results showed that sequence shuttles increase HDR efficiencies for all types of templates tested. We observed 11 ± 0.4%, 12 ± 0.3% and 10 ± 0.6% HDR efficiencies using 1 pmole of unstructured, looped or 18-helix nanostructures, respectively (Fig. 3B). To determine whether Cas9 RNPs bind directly to the DNA nanostructures as intended (*29*–*31*), we incubated the DNA with RNPs and used AFM to analyze the resulting samples. The images reveal Cas9 RNPs bound to the side of the 18-helix nanostructure, where the homology arms and the shuttle sequences are visibly located (Fig. 3C; Fig. S37, 38). We also investigated whether this strategy could be used to deliver nanostructured DNA into a different cell type. We electroporated synchronized human immortalized myelogenous leukemia K562 cells with Cas9 RNPs alongside unstructured, looped or 18-helix nanostructures including shuttle sequences. We observed similar HDR efficiencies with all templates, 5 ± 0.8%, 5 ± 0.5% and 4 ± 0.7% for the unstructured, looped or 18-helix nanostructures, respectively (Fig. 3D). Overall, these results demonstrated that shuttle sequences can increase the rate of nanostructured DNA incorporation into the genome in different cell lines and at similar levels relative to unstructured ssDNA templates.

**Figure 3:**
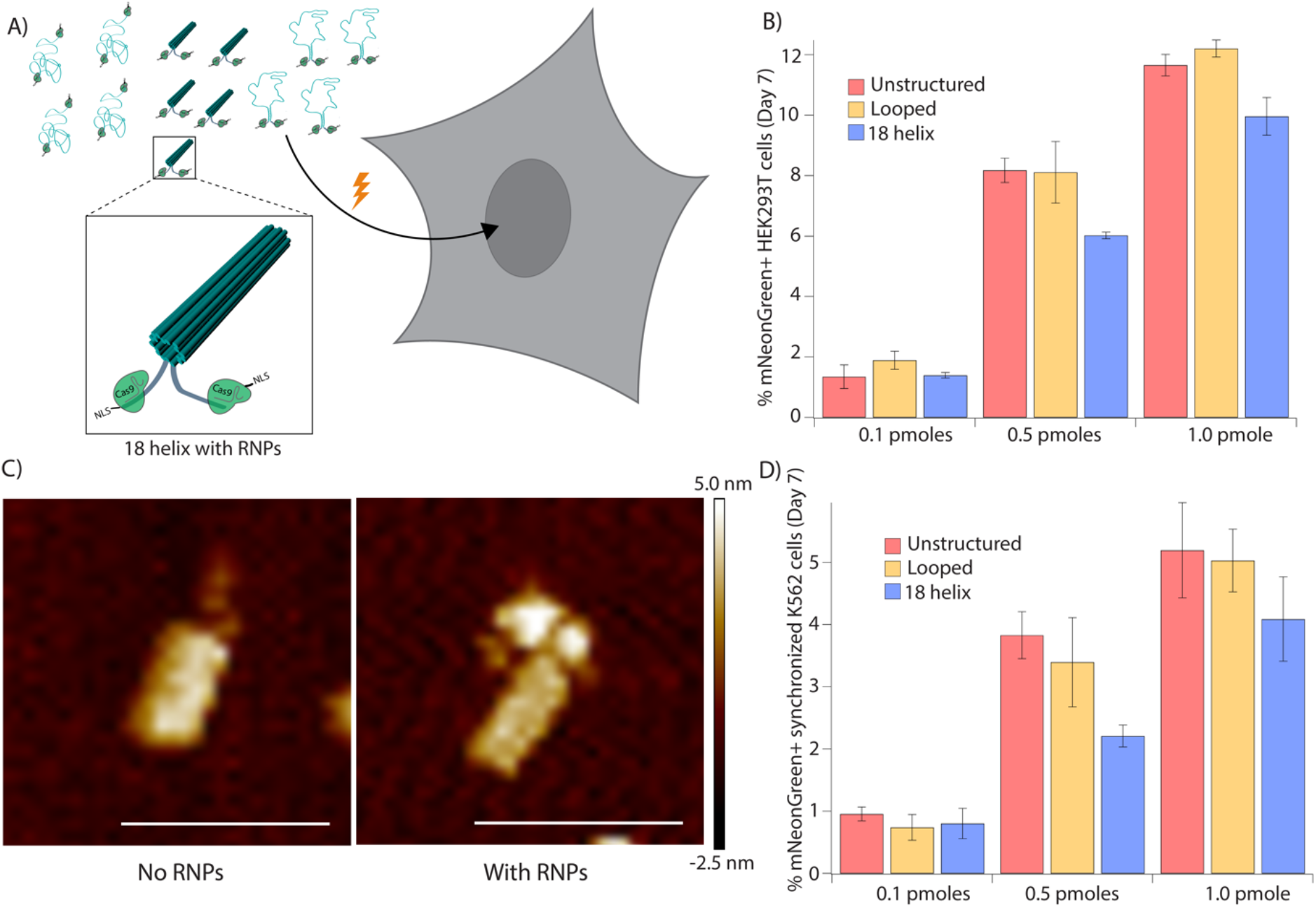
CRISPR-Cas9 RNP localization at template DNA ends increases HDR efficiency. **A)** Schematic of CRISPR-Cas9 binding to the ends of unstructured, looped and 18 helix nanostructure templates. CRISPR-Cas9 carries nuclear localization signals (NLS) to enter the nucleus upon electroporation. **B)** Aggregated flow cytometry data shows knock-in efficiencies are similar across unstructured, looped and 18 helix nanostructure templates when electroporating templates bound by CRISPR-Cas9 RNP. Error bars represent SDs from 3 experiments. **C)** AFM image depicting CRISPR-Cas9 RNPs i.unbound and ii. bound to 18 helix DNA nanostructures. Scale bar: 100 nm. **D)** Experiments in synchronized K562 cells show comparable knock-in efficiencies across unstructured, looped and 18 helix nanostructure. Error bars represent SDs from 3 experiments.

### Nanostructured DNA comprising a human gene enhances human primary cell HDR

We next investigated nanostructured DNA delivery into primary human T cells using a ~3.5 kb multigene cassette targeting *IL2RA*, a gene mutated in some families with a monogenic immune disorder that is potentially amenable to gene replacement strategies (OMIM 606367) (*32*–*34*). A ssDNA scaffold composed of two ~300 bp homology arms flanking the entire *IL2RA* open reading frame (ORF), fused to a GFP-encoding sequence and a separate mCherry-encoding sequence, was tested as an HDR template (Fig. 4A). Insertion of this HDR template into the genome results in co-expression of a detectable IL2RA-GFP fusion protein driven by the endogenous IL2RA promoter and a separate mCherry protein driven by the EF1a promoter. GFP expression would indicate truncated insertion, and expression of mCherry alone could indicate either truncation or insertion into an off-target genomic locus. Using a similar 18-helix nanostructure design as above, four different versions of these DNA nanostructures were created with either an alternating pattern of base pairing (50% Staples), 18-helix nanostructure restricted to the top half where the homology arms are located (Only Top), an open sheet-like structure (Open), and the full 18-helix nanostructure (Complex) (Fig. 4B) (Fig. S1-16). We also tested a looped structure comprising only five short oligonucleotide-directed helices, an unstructured ssDNA and an unstructured dsDNA for comparison. All HDR templates included shuttle sequences, and we used electroporation along with Cas9 RNPs on primary human T cells from two different human blood donors for this experiment. All DNA nanostructures demonstrated similar DNA insertion efficiencies compared to long unstructured ssDNA, consistent with our results in HEK293T and K526 cells (Fig. 3; Fig. 4C). In line with previous reports, ssDNA templates demonstrated lower toxicity and higher HDR efficiency relative to dsDNA controls (*35*, *36*) (Fig. 4C). Interestingly, although the DNA nanostructures comprised folded dsDNA segments, they caused less toxicity than the unstructured dsDNA templates tested here. A possible explanation is that the compact nature of the DNA nanostructures, where dsDNA helices are packed closely together and thus are less accessible, may circumvent mechanisms driving the toxicity of free dsDNA. Overall, these results demonstrated that complex DNA nanostructures can be used to compress large ssDNA HDR templates and can mediate efficient insertion in primary human T cells at endogenous target loci.

**Figure 4:**
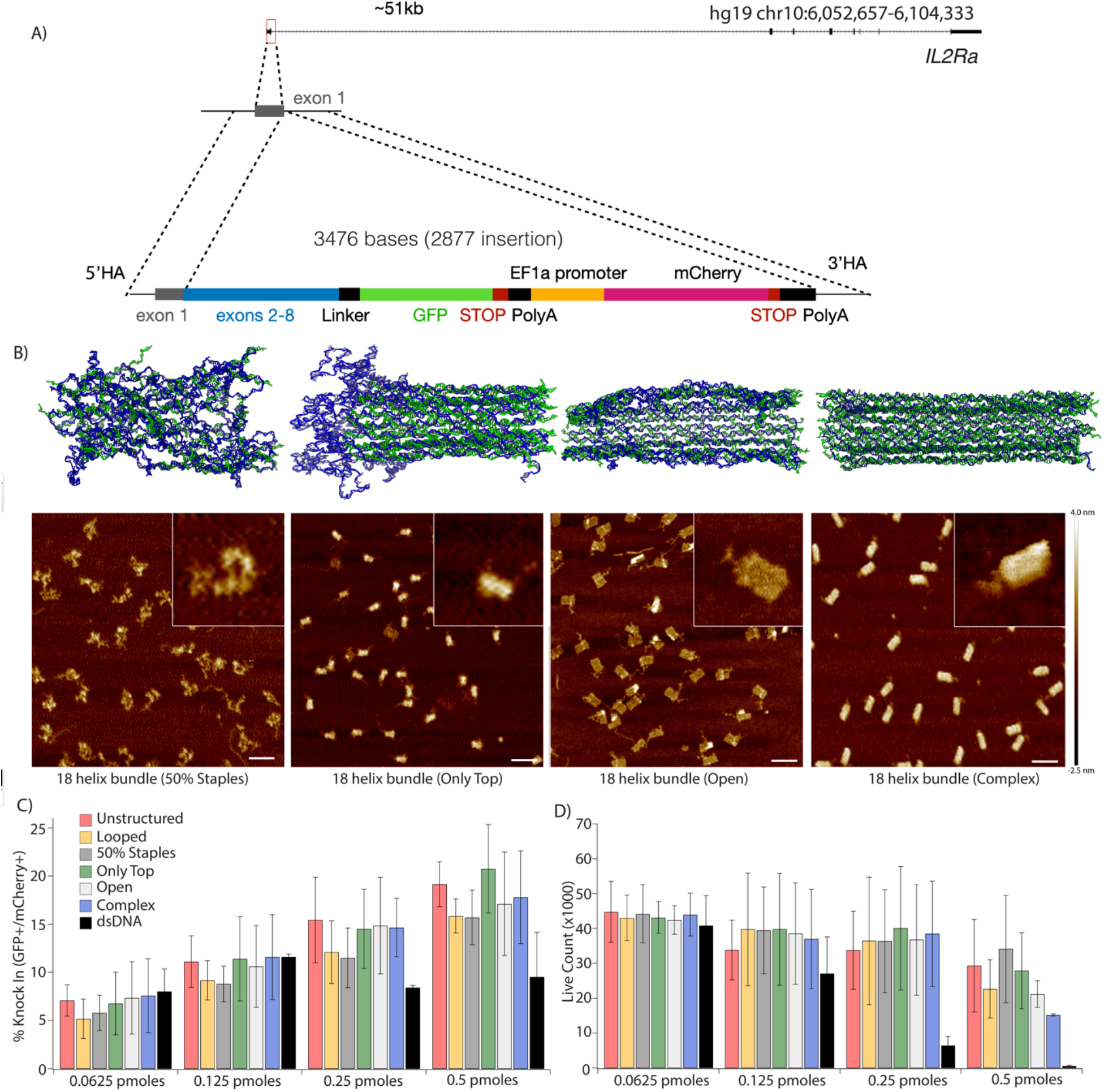
Nanostructured DNA comprising a human gene enhances human primary cell HDR. **A)** Schematic of knock-in strategy of a 3.5 kb HDR template encoding *IL2RA-*GFP fusion and mCherry driven by an EF1a promoter. **B)** oxDNA simulations and AFM images of 4 distinct versions of 18 helix DNA nanostructured HDR templates, including 50% Staples, Only Top, Open and Complex. Scale bar: 100 nm. **C)** Unstructured ssDNA and 18 helix nanostructure templates show increased knock-in efficiency compared to dsDNA. Error bars represent SDs from duplicate experiments. **D)** Live cell count shows unstructured ssDNA and 18 helix nanostructured templates display lower toxicity compared to dsDNA. Error bars represent SDs from duplicate experiments.

### Virus-like particles enable efficient intracellular delivery of nanostructured DNA

We investigated whether compaction of ssDNA HDR templates in the form of DNA nanostructures can improve their delivery into HEK293T cells using Cas9-VLPs (*37*). To this end, we delivered shuttled unstructured, looped and 18-helix nanostructure mNeonGreen HDR templates (Fig. 1B) into HEK293T cells either by electroporation or using Cas9-VLPs (Fig. 5A). On day 7 after delivery, we collected and analyzed the cells using flow cytometry to track mNeonGreen+ cells. Consistent with previous results, the unstructured and structured DNAs introduced by electroporation yielded similar HDR efficiencies of ~15% in each case. Although VLP delivery reduced overall HDR levels, we observed a 2.5-fold increase in HDR efficiency for the 18-helix nanostructure templates (from <2% to >5%) compared to unstructured and looped templates. These results show that nanostructured DNA can be delivered using VLPs, providing the possibility of *in vivo* delivery for therapeutic or bioimaging purposes into specific tissues. Furthermore, these data demonstrated higher HDR efficiencies when combining Cas9-VLPs and nanostructured HDR templates, an important step towards creating *in vivo* gene replacement or modification therapies.

**Figure 5:**
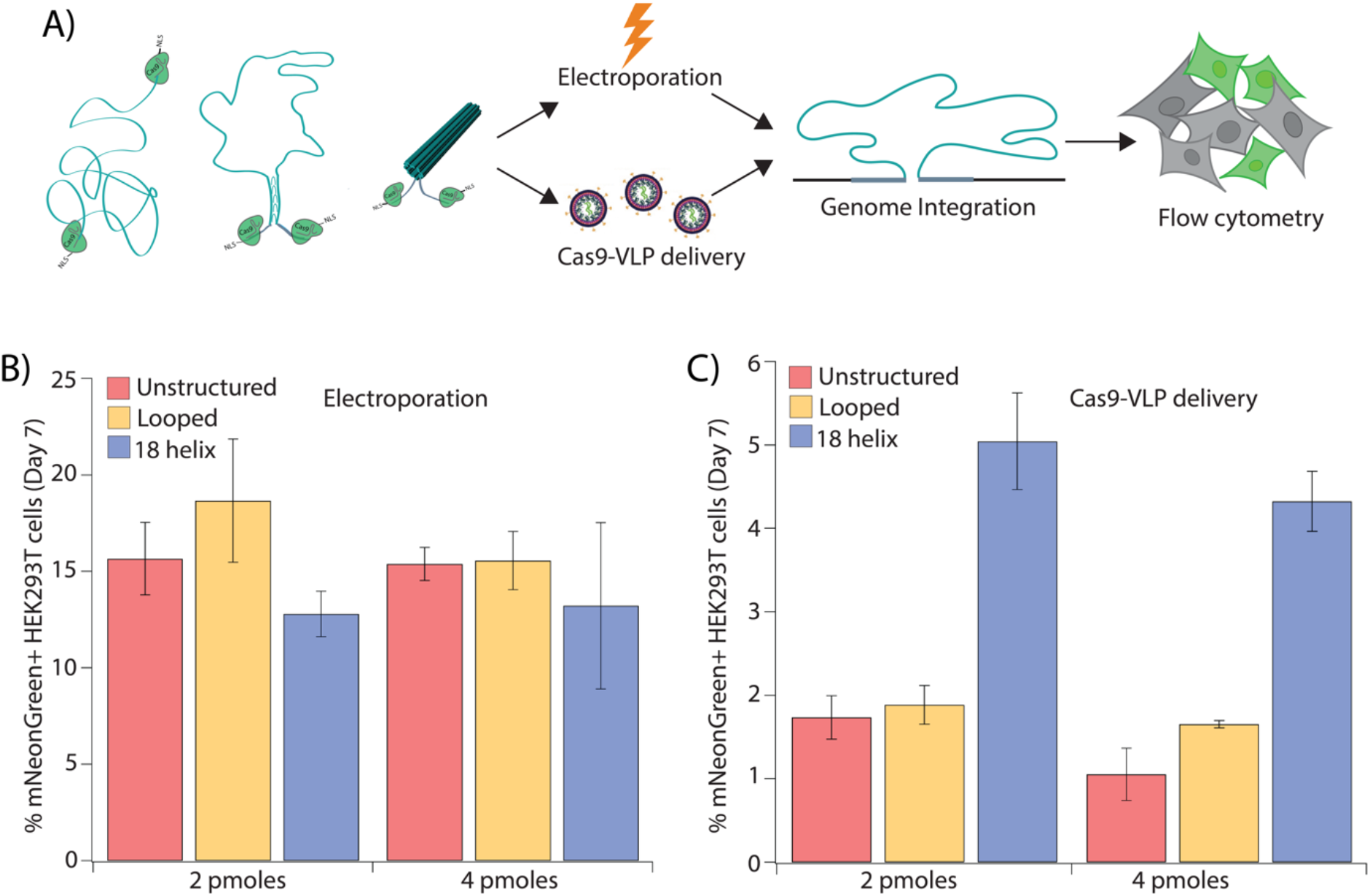
Virus-like particles enable efficient intracellular delivery of nanostructured DNA. **A)** Schematic of experimental setup where successful incorporation of HDR templates results in mNeonGreen+ cells. **B)** Knock-in efficiencies of unstructured, looped and 18 helix nanostructure show comparable values for delivery using electroporation. Error bars represent SDs from duplicate experiments. **C)** Cas9-VLP delivery shows 18 helix nanostructured templates display a 2.5-fold higher knock-in efficiency compared to unstructured and looped templates. Error bars represent SDs from duplicate experiments.

## Discussion

Programmed self-assembled DNA nanostructures have the ability to carry both engineered design features as well as genetic information, offering a route to creating novel therapeutic approaches and improving genome engineering methods. To date, versatile DNA nanostructures have been created, displaying three-dimensional structure, curvature, reconfiguration, modular design and hierarchical assembly into micrometer arrays (*5*, *38*, *39*). Despite these design advances, their ability to carry genetic information has been largely ignored. Nonetheless, potential *in cell* applications of DNA nanostructures have been discussed since their inception. In particular, *in vivo* use of DNA nanostructures for drug delivery was one of their first proposed applications (*40*). However, progress has been hampered by challenges, including cellular uptake and the stability of structures in cells (*11*). DNA nanostructures have demonstrated promise for the delivery of molecular payloads including small molecule drugs, siRNA, peptides, and proteins *in vitro* (*13*, *41*–*43*) and *in vivo* (*16*, *44*–*46*). These studies utilize the DNA nanostructure as a carrier, taking advantage of the ability to precisely incorporate these molecules on internal or external surfaces. Here we leveraged the ability to package a gene-length sequence, making the information encoded in the nanostructure itself the payload. In contrast to prior studies, gene delivery requires entry to cell nuclei, which has not been previously demonstrated. A recent study showed successful electroporation of DNA nanostructures into mammalian cells (*12*) but only demonstrated delivery into the cytosol.

Here we describe the use of nanostructured DNAs as templates for HDR-mediated genome editing using CRISPR-Cas9, providing a strategy for DNA compaction and localization that could expand CRISPR applications. This is the first instance of DNA nanostructures generated from scaffolds containing genes that could be delivered into human cells by electroporation or using Cas9-VLPs. We found that addition of truncated Cas9 target site sequences onto the ends of the nanostructured DNA improves HDR efficiency, presumably due to enhanced template localization to the site of genome repair following Cas9 cleavage (*27*, *28*). Furthermore, nanostructured DNA could be delivered into primary human T cells and used as donor templates for gene replacement following Cas9-catalyzed genome cleavage. Finally, we observed higher HDR efficiency for nanostructured versus unstructured DNA templates when delivered by VLPs, opening the possibility of introducing nanostructured DNA templates *in vivo* in a tissue-specific manner for therapeutic applications.

Truncated Cas9 target sequences can increase template delivery into the nucleus and increase HDR efficiency of DNA templates (*27*, *28*). AFM experiments showed that Cas9 associates with these sequences, which may induce enhanced template localization as passengers during nuclear import of Cas9 RNPs. Enhanced Cas9-induced HDR with tag-containing DNA templates was observed in multiple cell types including HEK293T, K562 and human primary T cells. Notably, although higher HDR levels and lower toxicity occurred with nanostructured DNA compared to unstructured dsDNA templates, we did not observe increased HDR efficiency compared to unstructured ssDNA templates. This suggests that HDR efficiency may be further increased by changing design features of the nanostructures, pending insight into the mechanisms underlying nuclear localization and genomic integration of structured DNA.

Nanostructured DNA delivery using Cas9-VLPs has two primary advantages over electroporation: lower toxicity and the potential for tissue-specific and *in vivo* delivery (*37*). Our data show a higher HDR efficiency when coupling Cas9-VLPs with DNA nanostructured templates, compared to unstructured ssDNA templates. How compaction of templates into nanostructures improves VLP delivery remains an open question. However, this strategy offers the potential to deliver DNA nanostructures *in vivo* in a tissue-specific manner, which could enhance bioimaging, radiotherapy and cancer treatment applications. It further allows *in vivo* delivery of large HDR DNA templates in diseases in which a gene replacement at the endogenous site could serve as a universal cure for patients suffering from a wide range of different substitution mutations and deletions on the causal genes.

Together, these findings validate three distinct strategies to deliver nanostructured DNA into cell nuclei and demonstrate their utility as templates for HDR-mediated genome editing. By exploiting their design features together with their capacity to carry genetic information, DNA nanostructures provide a new approach to DNA template-based genetic manipulation that could enable tissue-selective delivery and editing using Cas9-VLPs.

## Materials and Methods

### ssDNA production

Biotin-labeled dsDNA template was first amplified from the plasmid encoding the template design with biotin-labeled primers using KAPA HiFi HotStart ReadyMix PCR Kit (Roche) for 35 cycles (98°C for 20 s, 65°C for 15 s, and 72°C for 1 min; then 72°C for 1 min). dsDNA was then purified and concentrated by mixing with 1.8x sample volume of SPRI beads (UC Berkeley Sequencing Core). The samples were placed on a DynaMag-2 magnet (Thermo Fisher Scientific) for 5 min, and the supernatant was removed. The samples were washed twice with 70% ethanol and eluted in Tris-EDTA Buffer (Corning).

ssDNA was prepared by separating dsDNA strands using Dynabeads MyOne Streptavidin C1 beads (Thermo Fisher Scientific). Streptavidin beads were first washed three times with 1 mL 1x binding and washing (B&W) buffer (2x B&W buffer: 10 mM Tris-HCl pH 7.5, 1 mM EDTA, 2.0 M NaCl). dsDNA was added to Streptavidin beads and rotated for 30 min at room temperature. Beads were then collected on a magnet and washed once with 1x B&W buffer. Next, beads were resuspended in 2×100 uL melt buffer (125 mM NaOH in ddH2O), incubated for 2 min, and immediately precipitated, and the supernatant was transferred to a new non-stick 1.5 mL microcentrifuge tube containing 1 mL neutralization buffer (60 mM acetate in TE buffer pH 8.0). The supernatant containing ssDNA was purified using SPRI beads and eluted in ddH2O.

### Folding and purification of DNA nanostructures

DNA nanostructures were folded by mixing 10 nM ssDNA HDR template with 100 nM staple strands in 1x TEMg buffer (5 mM Tris, 1 mM EDTA, 8 mM MgCl_2_, pH 8). The thermal annealing protocol starts with a heat denaturation step at 65°C to remove any undesired secondary structure and then gradually decreases over the course of 14 hours to 20°C (table S1). DNA nanostructures were subsequently purified and concentrated by 5-6 rounds of spin filtration (Amicon 100kDa) at 5,000 rcf. Samples for AFM imaging were purified by using the Freeze ‘N Squeeze kit (BioRad) according to the manufacturer’s protocol.

### Agarose gel electrophoresis

Proper folding of DNA nanostructures was confirmed using agarose gel electrophoresis. 150 fmol of folded sample was loaded into agarose gels (1.5% agarose, 1x TBEMg (45 mM Tris, 45 mM boric acid, 1 mM EDTA, 11 mM MgCl_2_, pH 8), containing Ethidium bromide) and ran for 90 minutes at 90 volt submerged in an ice-water bath. Insertion of the mNeonGreen construct was confirmed using PCR amplification (PrimeSTAR® GXL DNA Polymerase, catalog R050B) of the target site followed by agarose gel electrophoresis. PCR products were loaded into agarose gels (2% agarose, 1x TAE, containing SYBR Safe) and ran for 90 minutes at 100 volt.

### Atomic force microscopy

Freeze ‘N Squeeze purified DNA nanostructure samples were imaged with a Bruker BioScope Resolve using the ScanAsyst in Air mode. Samples were prepared by applying 6 μl of sample to freshly cleaved mica (Plano GmbH) and 3 minutes of incubation before the mica was carefully rinsed with ddH2O and dried with a gentle flow of air. Imaging was performed with ScanAsyst-Air probes at a typical scan rate of around 1 Hz.

### oxDNA simulations

Simulations of four distinct versions of 18 helix DNA nanostructured HDR templates were performed using the coarse-grained model oxDNA (*26*), including 50% Staples, Only Top, Open and Complex. First, the original caDNAno (*47*) scaffold and staple strand routings of each nanostructure version were converted to the oxDNA model utilizing the tacoxDNA source code (http://tacoxdna.sissa.it/). Then, a multistep relaxation (table S2) was done to obtain an initial geometry for MD simulations. The relaxation steps are required to correct for overstretched bonds that result from the caDNAno to oxDNA conversion and to resemble a more realistic geometry. Once the structures were relaxed the MD simulations were performed: Consisting of 1 × 10^7 steps for the structures without homology arms and 1.1 to 2.5 × 10^8 steps for the ssDNAIn and structures containing homology arms with a time step of 0.005, which translates to 0.1 μs to 2.5 μs in real-time units. All simulations were performed without applying any external forces and implementing the oxDNA2 package and the NVE John thermostat at a temperature of 303 K and a salt concentration equivalent to 0.5 M NaCl. To run the simulations more efficiently GPU acceleration was implemented using the OSC (Ohio Supercomputer Center) resources. Analysis of the structural properties was performed using the software MagicDNA (*39*) and python-based analysis tool package (https://github.com/sulcgroup/oxdna_analysis_tools) (table S3; figs. S2, 3, 6, 7, 10, 11, 14, 15, 18, 19, 22, 23, 26, 27, 29, 30, 31-34; movies S1-4). Lastly, UCSF Chimera software was used for image and video rendering (Fig.1C; fig S1, 5, 9, 13, 17, 21, 25, 28) (*48*).

### Cell culture

HEK293T and K562 cells were cultured with 10% fetal bovine serum (VWR) and 1% Penicillin/Streptomycin (Gibco) at 37°C in a 5% CO2 air incubator. HEK293T cells were cultured in DMEM (Corning), and K562 cells were cultured in RPMI-1640 (Gibco) media. Routine checks for mycoplasma contamination were performed using the MycoAlert mycoplasma detection kit (Lonza).

### Transfections

Transfection was performed using Lipofectamine 3000 (Thermo Fisher Scientific) according to the manufacturer’s instructions. 50,000 cells per well were seeded in 24-well plates 24 hours prior to lipofection. Cells were transfected with 500 ng Cas nuclease expression plasmid, 150 ng sgRNA expression plasmid, and 0.5 pmol of either unstructured ssDNA, looped or 18 helix nanostructures per well.

### RNP Electroporation

Cas9 RNPs were formulated as previously described (*27*). crRNA and tracrRNA (Horizon Discovery) were resuspended in IDT duplex buffer with a polyglutamic acid or ssDNAenh electroporation enhancer (IDT) and stored in aliquots at −80°C until use. Immediately prior to electroporation, crRNA and tracrRNA were thawed and annealed at a 1:1 molar ratio to form gRNA. gRNA and Cas9-NLS (UC Berkeley QB3 MacroLab) were then mixed at a 2:1 molar ratio to form Cas9 RNPs. Electroporation was performed using a 96-well format 4D nucleofector (Lonza). HEK293T cells were electroporated with the SF buffer and the CM-130 pulse code, K562 cells with SF buffer and the FF-120 pulse code, and T cells with the P3 buffer and EH-115 pulse code. Cells were immediately resuspended in pre-warmed media, incubated for 20 minutes, and transferred to culture plates.

### Flow cytometry

Primary human T cells were collected 5 days after electroporations, resuspended in FACS buffer, and stained with GhostDye red 780 (Tonbo), anti-human CD4-PerCP (Tonbo, Cat #67-0047-T500), and anti-human CD25-BV421 (Biolegend, Cat #302630). All primary human T cell gating strategies included singlet gating, live:dead differentiation, and CD4 and CD8 T cell differentiation and excluded subcellular debris. All quantified data for experiments using primary human T cells were collected from gated CD4+ T cells only.

Cells were resuspended in FACS buffer (1% BSA in PBS) and analyzed by flow cytometry for mNeonGreen positive cells 7 days post-transfection. Flow cytometry was performed on an Attune NxT flow cytometer with a 96-well autosampler (Thermo Fisher Scientific), and data analysis was performed using the FlowJo v10 software.

### Primary Human T cell culture

Leukapheresis products from anonymous healthy human donors were purchased from STEMCELL Technologies, Inc. and isolated using an EasySep human T cell isolation kit (Cat #17951). Isolated CD3+ T cells were activated at 1×10^6^ cells mL^−1^ in a 1:1 ratio with CD3/CD28 magnetic dynabeads (CTS, Thermo Fisher Scientific), 100 U mL^−1^ of IL-7, and 10 U mL^−1^ IL-15 (R&D Systems) for 48 hours in complete XVivo15 medium (Lonza) (5% fetal bovine serum, 50 μM 2-mercaptoethanol, 10 mM N-alcetyl L-cysteine). Following the 48-hour activation period, CD3+ T cells were debeaded with an EasySep magnet (STEMCELL) prior to resuspension and electroporation at 0.5-1×10^6^ cell mL^−1^ in P3 buffer (Lonza). Following electroporations with RNPs and HDRTs, fresh medium and cytokines were added every 2-3 days.

### Cas9-VLP

Cas9-VLPs were harvested from transfected Lenti-X cells. Cultured cells were transfected with 1 μg VSV-G, 3.3 μg psPax2, 6.7 μg Gag-Cas9, and 10 μg U6-ELS77 plasmids using polyethylenimine (Polyscience Inc.). Transfected cells were switched into Optimem (Gibco) 12 hours post transfection and supernatants were harvested 48 hours post media change. Supernatants were pooled, filtered through a .45um aPES filter bottle (Thermo Fisher Scientific). Filtered samples were then concentrated via ultracentrifugation for 2 hours at 25,000rpm on a 30% sucrose cushion. Concentrated VLPs were resuspended in SE buffer (Lonza) and electroporated with 2-4pmol HDR template using a 4-D Nucleofector with pulse code CM-150 (Lonza). VLP/template mix was added to 15,000 cells in 50ul DMEM +10% FBS and 1x penicillin/streptomycin.

Following a 30 minute incubation at 37C, 75 ul Optimem was added to bring the final well volume to approximately 150 ul. Cells were passaged on day 3 to maintain sub-confluent culture conditions and analyzed by flow cytometry on day 7 using an Attune NxT (Thermo Fisher Scientific).

## Author contributions

This project was conceived by E.L.-S. and J.A.D. E.L.-S. initiated and led the study with input from W.G.P., C.E.C., M.G.P., R.L., B.R.S. and A.M. W.G.P. designed and characterized the DNA nanostructures with input from C.E.C., M.G.P, D.M.L. and E.L.-S. D.M.L. performed structure simulations on oxDNA with input from W.G.P and C.E.C. E.L.-S., M.S.D., A.D. and E.C. purified, folded and concentrated DNA nanostructures for *in vivo* experiments with input from W.G.P. B.R.S. and V.S.V. performed experiments in human primary T cells. J.R.H. and C.R.S.E. designed and performed VLP experiments with input from E.L.-S. E.L.-S. and E.C.S. designed and performed further in cell experiments. E.L.-S., W.G.P. and J.A.D. wrote the manuscript. All authors reviewed and commented on the manuscript.

## Data and materials availability

All data needed to evaluate the conclusions in the paper are present in the paper and/or the Supplementary Materials.

## List of Supplementary Materials

Figure S1: oxDNA relaxation of the 18 helix nanostructure (complex; GFP/mCherry template).

Figure S2: Root-mean-square fluctuation analysis of oxDNA trajectories of the 18 helix nanostructure (complex; GFP/mCherry template).

Figure S3: Root-mean-square deviation analysis of oxDNA trajectories of the 18 helix nanostructure (complex; GFP/mCherry template).

Figure S4: Representative AFM image of the 18 helix structure (complex; GFP/mCherry template).

Figure S5: oxDNA relaxation of the only top structure (GFP/mCherry template).

Figure S6: Root-mean-square fluctuation analysis of oxDNA trajectories of the only top nanostructure (GFP/mCherry template).

Figure S7: Root-mean-square deviation analysis of oxDNA trajectories of the only top nanostructure (GFP/mCherry template).

Figure S8: Representative AFM image of the only top nanostructure (GFP/mCherry template).

Figure S9: oxDNA relaxation of the open nanostructure (GFP/mCherry template).

Figure S10: Root-mean-square fluctuation analysis of oxDNA trajectories of the open nanostructure (GFP/mCherry template).

Figure S11: Root-mean-square deviation analysis of oxDNA trajectories of the open nanostructure (GFP/mCherry template).

Figure S12: Representative AFM image of the open nanostructure (GFP/mCherry template).

Figure S13: oxDNA relaxation of the 50% staples nanostructure (GFP/mCherry template).

Figure S14: Root-mean-square fluctuation analysis of oxDNA trajectories of the 50% staples nanostructure (GFP/mCherry template).

Figure S15: Root-mean-square deviation analysis of oxDNA trajectories of the 50% staples nanostructure (GFP/mCherry template).

Figure S16: Representative AFM image of the 50% staples nanostructure (GFP/mCherry template).

Figure S17: oxDNA relaxation of the 18 helix nanostructure (complex), including ssDNA homology arms (mNeon template).

Figure S18: Root-mean-square fluctuation analysis of oxDNA trajectories of the 18 helix nanostructure (complex), including ssDNA homology arms (mNeon template).

Figure S19: Root-mean-square deviation analysis of oxDNA trajectories of the 18 helix nanostructure (complex), including ssDNA homology arms (mNeon template).

Figure S20: Representative AFM image of the 18 helix nanostructure (complex) (mNeon template).

Figure S21: oxDNA relaxation of the unstructured ssDNA (GFP/mCherry template).

Figure S22: Root-mean-square fluctuation analysis of oxDNA trajectories of the unstructured ssDNA (GFP/mCherry template).

Figure S23: Root-mean-square deviation analysis of oxDNA trajectories of the unstructured ssDNA (GFP/mCherry template).

Figure S24: Representative AFM image of the unstructured ssDNA (GFP/mCherry template).

Figure S25: oxDNA relaxation of the 18 helix nanostructure (complex), including ssDNA homology arms (GFP/mCherry template).

Figure S26: Root-mean-square fluctuation analysis of oxDNA trajectories of the 18 helix nanostructure (complex), including ssDNA homology arms (GFP/mCherry template).

Figure S27: Root-mean-square deviation analysis of oxDNA trajectories of the 18 helix nanostructure (complex), including ssDNA homology arms (GFP/mCherry template).

Figure S28: oxDNA relaxation of the unstructured ssDNA (mNeon template).

Figure S29: Root-mean-square fluctuation analysis of oxDNA trajectories of the unstructured ssDNA (mNeon template).

Figure S30: Root-mean-square deviation analysis of oxDNA trajectories of the unstructured ssDNA (mNeon template).

Figure S31: Distances between the shuttle sites on the homology arms of the unstructured ssDNA (GFP/mCherry template) by oxDNA.

Figure S32: Distances between the shuttle sites on the homology arms of the 18 helix nanostructure (complex; GFP/mCherry template) by oxDNA.

Figure S33: Distances between the shuttle sites on the homology arms of the unstructured ssDNA (mNeon template) by oxDNA.

Figure S34: Distances between the shuttle sites on the homology arms of the 18 helix nanostructure (complex; mNeon template) by oxDNA.

Figure S35: Agarose gel electrophoresis of 18 helix nanostructures on both templates.

Figure S36: Agarose gel electrophoresis of folded structures on the GFP/mCherry template.

Figure S37: Agarose gel electrophoresis of the 18 helix structure (GFP/mCherry) with RNPs.

Figure S38: Representative AFM image of the 18 helix nanostructure with RNPs (GFP/mCherry template).

Table S1. Thermal annealing protocol.

Table S2. oxDNA simulation parameters.

Table S3: Average distance between shuttle sites for 18 helix nanostructures and unstructured ssDNA on both HDR templates.

Movie S1. oxDNA model of the 18 helix nanostructure (GFP/mCherry template).

Movie S2. oxDNA model of the only Top nanostructure (GFP/mCherry template).

Movie S3. oxDNA model of the open nanostructure (GFP/mCherry template).

Movie S4. oxDNA model of the 50% staples nanostructure (GFP/mCherry template).

Movie S5. oxDNA MD simulation trajectory of the 18 helix tube with ssDNA homology arms (GFP/mCherry template).

Movie S6. oxDNA MD simulation trajectory of the unstructured ssDNA (GFP/mCherry template).

Movie S7. oxDNA MD simulation trajectory of the 18 helix tube with ssDNA homology arms (mNeon template)

Movie S8. oxDNA MD simulation trajectory of the unstructured ssDNA (mNeon).

Data file S1. caDNAno design files and staple strand sequences.

Data file S2. oxDNA simulation input files.

## References

1. N. C. Seeman, Nucleic acid junctions and lattices. J Theor Biol. 99, 237–247 (1982).

2. J. H. Chen, N. C. Seeman, Synthesis from DNA of a molecule with the connectivity of a cube. Nature. 350, 631–633 (1991).

3. P. W. K. Rothemund, Folding DNA to create nanoscale shapes and patterns. Nature. 440, 297–302 (2006).

4. C. E. Castro, F. Kilchherr, D.-N. Kim, E. L. Shiao, T. Wauer, P. Wortmann, M. Bathe, H. Dietz, A primer to scaffolded DNA origami. Nat Methods. 8, 221–229 (2011).

5. S. Dey, C. Fan, K. V. Gothelf, J. Li, C. Lin, L. Liu, N. Liu, M. A. D. Nijenhuis, B. Saccà, F. C. Simmel, H. Yan, P. Zhan, DNA origami. Nat Rev Methods Primers. 1, 1–24 (2021).

6. Y. Zhao, X. Dai, F. Wang, X. Zhang, C. Fan, X. Liu, Nanofabrication based on DNA nanotechnology. Nano Today. 26, 123–148 (2019).

7. D. Daems, W. Pfeifer, I. Rutten, B. Saccà, D. Spasic, J. Lammertyn, Three-dimensional DNA origami as programmable anchoring points for bioreceptors in fiber optic surface plasmon resonance biosensing. ACS Appl. Mater. Interfaces. 10, 23539–23547 (2018).

8. J. J. Funke, P. Ketterer, C. Lieleg, P. Korber, H. Dietz, Exploring nucleosome unwrapping using DNA origami. Nano Lett. 16, 7891–7898 (2016).

9. J. V. Le, Y. Luo, M. A. Darcy, C. R. Lucas, M. F. Goodwin, M. G. Poirier, C. E. Castro, Probing nucleosome stability with a DNA origami nanocaliper. ACS Nano. 10, 7073–7084 (2016).

10. Y. Wang, J. V. Le, K. Crocker, M. A. Darcy, P. D. Halley, D. Zhao, N. Andrioff, C. Croy, M. G. Poirier, R. Bundschuh, C. E. Castro, A nanoscale DNA force spectrometer capable of applying tension and compression on biomolecules. Nucleic Acids Research. 49, 8987–8999 (2021).

11. S. Ramakrishnan, H. Ijäs, V. Linko, A. Keller, Structural stability of DNA origami nanostructures under application-specific conditions. Computational and Structural Biotechnology Journal. 16, 342–349 (2018).

12. A. Chopra, S. Krishnan, F. C. Simmel, Electrotransfection of polyamine folded DNA origami structures. Nano Lett. 16, 6683–6690 (2016).

13. J. Li, C. Fan, H. Pei, J. Shi, Q. Huang, Smart drug delivery nanocarriers with self-assembled DNA nanostructures. Advanced Materials. 25, 4386–4396 (2013).

14. B. R. Madhanagopal, S. Zhang, E. Demirel, H. Wady, A. R. Chandrasekaran, DNA nanocarriers: programmed to deliver. Trends Biochem Sci. 43, 997–1013 (2018).

15. S. Zhao, F. Duan, S. Liu, T. Wu, Y. Shang, R. Tian, J. Liu, Z.-G. Wang, Q. Jiang, B. Ding, Efficient intracellular delivery of RNase A using DNA origami carriers. ACS Appl Mater Interfaces. 11, 11112–11118 (2019).

16. S. Li, Q. Jiang, S. Liu, Y. Zhang, Y. Tian, C. Song, J. Wang, Y. Zou, G. J. Anderson, J.-Y. Han, Y. Chang, Y. Liu, C. Zhang, L. Chen, G. Zhou, G. Nie, H. Yan, B. Ding, Y. Zhao, A DNA nanorobot functions as a cancer therapeutic in response to a molecular trigger in vivo. Nat Biotechnol. 36, 258–264 (2018).

17. Z. Wang, L. Song, Q. Liu, R. Tian, Y. Shang, F. Liu, S. Liu, S. Zhao, Z. Han, J. Sun, Q. Jiang, B. Ding, A tubular DNA nanodevice as a siRNA/chemo-drug co-delivery vehicle for combined cancer therapy. Angew Chem Int Ed Engl. 60, 2594–2598 (2021).

18. H. Zhang, G. S. Demirer, H. Zhang, T. Ye, N. S. Goh, A. J. Aditham, F. J. Cunningham, C. Fan, M. P. Landry, DNA nanostructures coordinate gene silencing in mature plants. PNAS. 116, 7543–7548 (2019).

19. S. M. Beltrán, M. J. Slepian, R. E. Taylor, Extending the capabilities of molecular force sensors via DNA nanotechnology. Crit Rev Biomed Eng. 48, 1–16 (2020).

20. S. Wang, Z. Zhou, N. Ma, S. Yang, K. Li, C. Teng, Y. Ke, Y. Tian, DNA origami-enabled biosensors. Sensors (Basel). 20, 6899 (2020).

21. E.-C. Wamhoff, J. L. Banal, W. P. Bricker, T. R. Shepherd, M. F. Parsons, R. Veneziano, M. B. Stone, H. Jun, X. Wang, M. Bathe, Programming structured DNA assemblies to probe biophysical processes. Annual Review of Biophysics. 48, 395–419 (2019).

22. N. C. Shaner, G. G. Lambert, A. Chammas, Y. Ni, P. J. Cranfill, M. A. Baird, B. R. Sell, J. Allen, R. N. Day, M. Israelsson, M. W. Davidson, J. Wang, A bright monomeric green fluorescent protein derived from Branchiostoma lanceolatum. Nat Methods. 10, 407–409 (2013).

23. Y. Ke, G. Bellot, N. V. Voigt, E. Fradkov, W. M. Shih, Two design strategies for enhancement of multilayer–DNA-origami folding: underwinding for specific intercalator rescue and staple-break positioning. Chem. Sci. 3, 2587–2597 (2012).

24. T. G. Martin, H. Dietz, Magnesium-free self-assembly of multi-layer DNA objects. Nat Commun. 3, 1103 (2012).

25. J. Y. Lee, J. G. Lee, G. Yun, C. Lee, Y.-J. Kim, K. S. Kim, T. H. Kim, D.-N. Kim, Rapid computational analysis of DNA origami assemblies at near-atomic resolution. ACS Nano. 15, 1002–1015 (2021).

26. P. Šulc, F. Romano, T. E. Ouldridge, L. Rovigatti, J. P. K. Doye, A. A. Louis, Sequence-dependent thermodynamics of a coarse-grained DNA model. J Chem Phys. 137, 135101 (2012).

27. D. N. Nguyen, T. L. Roth, P. J. Li, P. A. Chen, R. Apathy, M. R. Mamedov, L. T. Vo, V. R. Tobin, D. Goodman, E. Shifrut, J. A. Bluestone, J. M. Puck, F. C. Szoka, A. Marson, Polymer-stabilized Cas9 nanoparticles and modified repair templates increase genome editing efficiency. Nat Biotechnol. 38, 44–49 (2020).

28. B. R. Shy, V. Vykunta, A. Ha, T. L. Roth, A. Talbot, D. N. Nguyen, Y. Y. Chen, F. Blaeschke, S. Vedova, M. R. Mamedov, J.-Y. Chung, H. Li, J. Wolf, T. G. Martin, L. Ye, J. Eyquem, J. H. Esensten, A. Marson, in press, doi:10.1101/2021.09.02.458799.

29. M. Jinek, K. Chylinski, I. Fonfara, M. Hauer, J. A. Doudna, E. Charpentier, A programmable dual-RNA-guided DNA endonuclease in adaptive bacterial immunity. Science. 337, 816–821 (2012).

30. V. Pattanayak, S. Lin, J. P. Guilinger, E. Ma, J. A. Doudna, D. R. Liu, High-throughput profiling of off-target DNA cleavage reveals RNA-programmed Cas9 nuclease specificity. Nat Biotechnol. 31, 839–843 (2013).

31. J. K. Nuñez, L. B. Harrington, J. A. Doudna, Chemical and biophysical modulation of Cas9 for tunable genome engineering. ACS Chem Biol. 11, 681–688 (2016).

32. K. Goudy, D. Aydin, F. Barzaghi, E. Gambineri, M. Vignoli, S. Ciullini Mannurita, C. Doglioni, M. Ponzoni, M. P. Cicalese, A. Assanelli, A. Tommasini, I. Brigida, R. M. Dellepiane, S. Martino, S. Olek, A. Aiuti, F. Ciceri, M. G. Roncarolo, R. Bacchetta, Human IL2RA null mutation mediates immunodeficiency with lymphoproliferation and autoimmunity. Clin Immunol. 146, 248–261 (2013).

33. L. Bezrodnik, M. S. Caldirola, A. G. Seminario, I. Moreira, M. I. Gaillard, Follicular bronchiolitis as phenotype associated with CD25 deficiency. Clinical & Experimental Immunology. 175, 227–234 (2014).

34. A. Bousfiha, L. Jeddane, C. Picard, W. Al-Herz, F. Ailal, T. Chatila, C. Cunningham-Rundles, A. Etzioni, J. L. Franco, S. M. Holland, C. Klein, T. Morio, H. D. Ochs, E. Oksenhendler, J. Puck, T. R. Torgerson, J.-L. Casanova, K. E. Sullivan, S. G. Tangye, Human inborn errors of immunity: 2019 update of the IUIS phenotypical classification. J Clin Immunol. 40, 66–81 (2020).

35. V. Hornung, E. Latz, Intracellular DNA recognition. Nat Rev Immunol. 10, 123–130 (2010).

36. T. L. Roth, C. Puig-Saus, R. Yu, E. Shifrut, J. Carnevale, P. J. Li, J. Hiatt, J. Saco, P. Krystofinski, H. Li, V. Tobin, D. N. Nguyen, M. R. Lee, A. L. Putnam, A. L. Ferris, J. W. Chen, J.-N. Schickel, L. Pellerin, D. Carmody, G. Alkorta-Aranburu, D. del Gaudio, H. Matsumoto, M. Morell, Y. Mao, M. Cho, R. M. Quadros, C. B. Gurumurthy, B. Smith, M. Haugwitz, S. H. Hughes, J. S. Weissman, K. Schumann, J. H. Esensten, A. P. May, A. Ashworth, G. M. Kupfer, S. A. W. Greeley, R. Bacchetta, E. Meffre, M. G. Roncarolo, N. Romberg, K. C. Herold, A. Ribas, M. D. Leonetti, A. Marson, Reprogramming human T cell function and specificity with non-viral genome targeting. Nature. 559, 405–409 (2018).

37. J. R. Hamilton, C. A. Tsuchida, D. N. Nguyen, B. R. Shy, E. R. McGarrigle, C. R. Sandoval Espinoza, D. Carr, F. Blaeschke, A. Marson, J. A. Doudna, Targeted delivery of CRISPR-Cas9 and transgenes enables complex immune cell engineering. Cell Rep. 35, 109207 (2021).

38. M. DeLuca, Z. Shi, C. E. Castro, G. Arya, Dynamic DNA nanotechnology: toward functional nanoscale devices. Nanoscale Horiz. 5, 182–201 (2020).

39. C.-M. Huang, A. Kucinic, J. A. Johnson, H.-J. Su, C. E. Castro, Integrated computer-aided engineering and design for DNA assemblies. Nat. Mater. 20, 1264–1271 (2021).

40. S. M. Douglas, I. Bachelet, G. M. Church, A logic-gated nanorobot for targeted transport of molecular payloads. Science. 335, 831–834 (2012).

41. Q. Jiang, C. Song, J. Nangreave, X. Liu, L. Lin, D. Qiu, Z.-G. Wang, G. Zou, X. Liang, H. Yan, B. Ding, DNA origami as a carrier for circumvention of drug resistance. J Am Chem Soc. 134, 13396–13403 (2012).

42. Y.-X. Zhao, A. Shaw, X. Zeng, E. Benson, A. M. Nyström, B. Högberg, DNA origami delivery system for cancer therapy with tunable release properties. ACS Nano. 6, 8684–8691 (2012).

43. P. D. Halley, C. R. Lucas, E. M. McWilliams, M. J. Webber, R. A. Patton, C. Kural, D. M. Lucas, J. C. Byrd, C. E. Castro, Daunorubicin-loaded DNA origami nanostructures circumvent drug-resistance mechanisms in a leukemia model. Small. 12, 308–320 (2016).

44. S. Zhao, R. Tian, J. Wu, S. Liu, Y. Wang, M. Wen, Y. Shang, Q. Liu, Y. Li, Y. Guo, Z. Wang, T. Wang, Y. Zhao, H. Zhao, H. Cao, Y. Su, J. Sun, Q. Jiang, B. Ding, A DNA origami-based aptamer nanoarray for potent and reversible anticoagulation in hemodialysis. Nat Commun. 12, 358 (2021).

45. S. Palazzolo, M. Hadla, C. Russo Spena, I. Caligiuri, R. Rotondo, M. Adeel, V. Kumar, G. Corona, V. Canzonieri, G. Toffoli, F. Rizzolio, An effective multi-stage liposomal DNA origami nanosystem for in vivo cancer therapy. Cancers. 11, 1997 (2019).

46. Q. Zhang, Q. Jiang, N. Li, L. Dai, Q. Liu, L. Song, J. Wang, Y. Li, J. Tian, B. Ding, Y. Du, DNA origami as an in vivo drug delivery vehicle for cancer therapy. ACS Nano. 8, 6633–6643 (2014).

47. S. M. Douglas, A. H. Marblestone, S. Teerapittayanon, A. Vazquez, G. M. Church, W. M. Shih, Rapid prototyping of 3D DNA-origami shapes with caDNAno. Nucleic Acids Research. 37, 5001–5006 (2009).

48. E. F. Pettersen, T. D. Goddard, C. C. Huang, G. S. Couch, D. M. Greenblatt, E. C. Meng, T. E. Ferrin, UCSF Chimera--a visualization system for exploratory research and analysis. J Comput Chem. 25, 1605–1612 (2004).

49. C.-M. Huang, A. Kucinic, J. V. Le, C. E. Castro, H.-J. Su, Uncertainty quantification of a DNA origami mechanism using a coarse-grained model and kinematic variance analysis. Nanoscale. 11, 1647–1660 (2019).

50. E. Poppleton, J. Bohlin, M. Matthies, S. Sharma, F. Zhang, P. Šulc, Design, optimization and analysis of large DNA and RNA nanostructures through interactive visualization, editing and molecular simulation. Nucleic Acids Research. 48, e72 (2020).

