## Supplemental Information for "CRISPR-Cas9 mediated nuclear transport and genomic integration of nanostructured genes in human primary cells"

DNA origami material and folding:

Staple strands were purchased from IDT (www.idtdna.com) and Eurofins (www.eurofinsgenomics.com) as desalted products at 100 µM concentration in RNAse free water and used without further purification.

The protocol used for thermal annealing is as follows:

**Table S1: Thermal annealing protocol**

| T [°C] | min / °C |
|---|---|
| 65 | 15 |
| 64-61 | 3 |
| 60 | 5 |
| 59-58 | 10 |
| 57 | 15 |
| 56 | 25 |
| 55 | 30 |
| 54 | 45 |
| 53-49 | 60 |
| 48-45 | 42 |
| 44 | 36 |
| 43-42 | 32 |
| 41-39 | 20 |
| 38 | 15 |
| 37 | 10 |
| 36-35 | 5 |
| 34-30 | 2 |
| 20 | stay |

oxDNA simulations

**Table S2: oxDNA simulation parameters:** Simulation Input files available in supplemental information “data_file_S2.zip”

| Input | Steps | Dt (unit: 3.03 ps) | Relaxation Strength | Force in base paired bases | oxDNA2 | Use GPU |
|---|---|---|---|---|---|---|
| Relax_a | 1.5 x 10^4 | 10^-7 | 150 | Y | Y | N |
| Relax_b | 10^4 | 5 x 10^-5 | 150 | Y | Y | N |
| Relax_c | 2 x 10^4 | 5 x 10^-3 | 150 | Y | Y | N |
| Short Run | 10^6 | 5 x 10^-4 | N/A | Y | Y | Y |
| OxDNA | 10^7 ,10^8 or 2.510^8 | 5 x 10^-3 | N/A | N | Y | Y |

The relaxation (relax_a-c and short run) and simulations (input oxDNA) use oxDNA2, whose main upgrades include correction of structural parameters such as twist and the incorporation of minor and major grooves. During the relaxation steps it is possible for bases to move too much and cause simulation to fail or to  be unstable, to avoid such, the spring constants and the time steps are gradually increased. Once the relaxation steps a-c are completed a short simulation is run adding mutual forces between bases which would aid to maintain an equilibrium distance between parts of the structures in the cases in which the first relaxation steps cause large shifts (*49*). These forces are generated by selecting all the bases and importing a force file from oxView (*50*). Lastly an MD simulation is run  increasing the time steps to 5 x 10^-3 and ensuring that the aforementioned external forces are removed.

We then use magicDNA to analyze the simulation results and get the Root-mean-square fluctuation (RMSF) as a measure for the local fluctuations from the average configuration of each base over time. Additionally, we use magicDNA to get the Root-mean-square deviation (RMSD) to assess the average spatial deviation for each base, based on the initial configuration over time. We provide the average configuration over the simulated time steps as graphical models, with color-coded RMSF values for each structure. *N.B.* the upper and lower limit for the color-code are structure specific and provide visual support. For the correct comparison between different structures, use the corresponding numerical values.

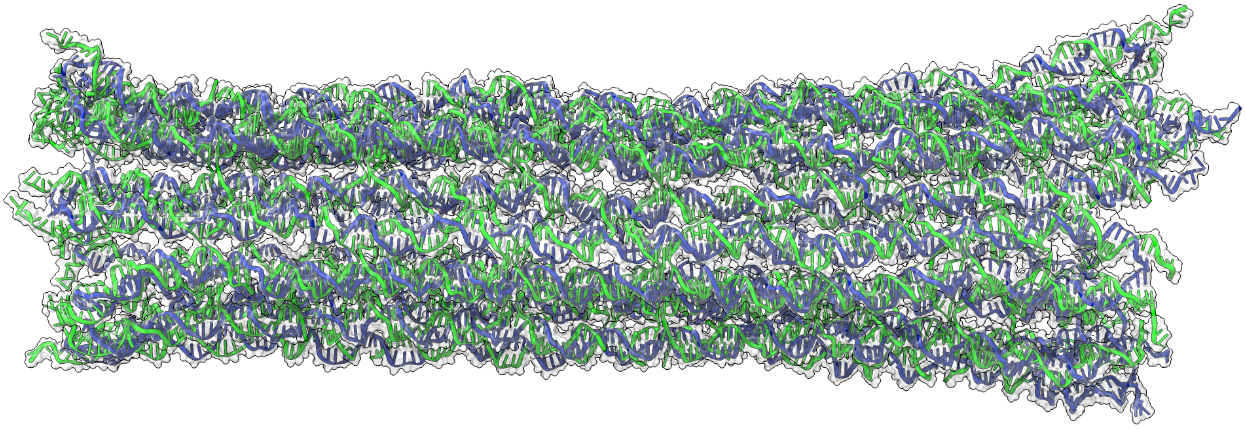

**Figure S1: oxDNA relaxation of the 18 helix nanostructure (complex; GFP/mCherry template).**
Green color represents staple strands, blue color represents the template DNA. Simulations were run for 10^7 steps. ssDNA homology arms are not included to improve visualization of the structure.

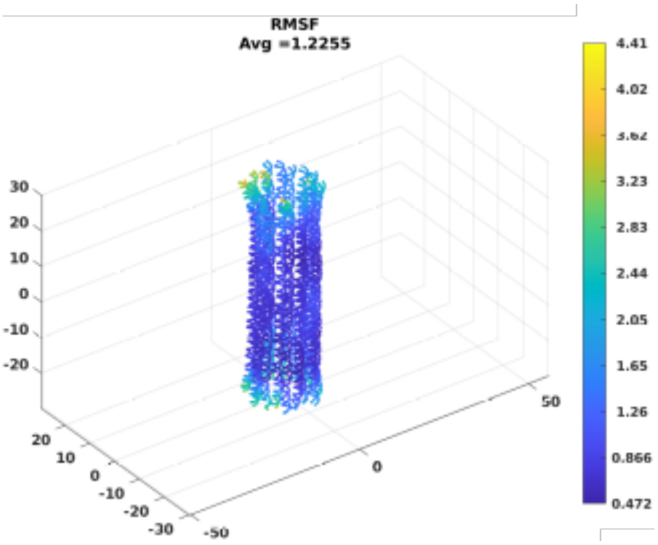


**Figure S2: Root-mean-square fluctuation analysis of oxDNA trajectories of the 18 helix nanostructure (complex; GFP/mCherry template).**
RMSF (in nm) of the 18 helix nanostructure through 10^7 steps in oxDNA. Dark blue areas indicate smaller fluctuations than yellow areas. Processed in magicDNA.

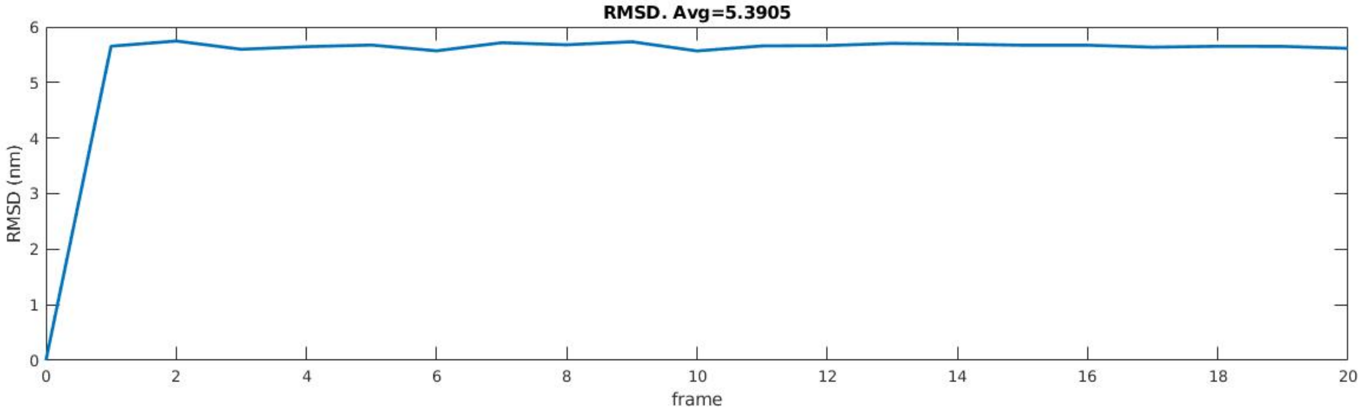


**Figure S3: Root-mean-square deviation analysis of oxDNA trajectories of the 18 helix nanostructure (complex; GFP/mCherry template).**

RMSD (in nm) of the 18 helix nanostructure through 10^7 steps in oxDNA. The graph indicates steady-state behavior as the RMSD reaches its plateau. Processed in magicDNA.

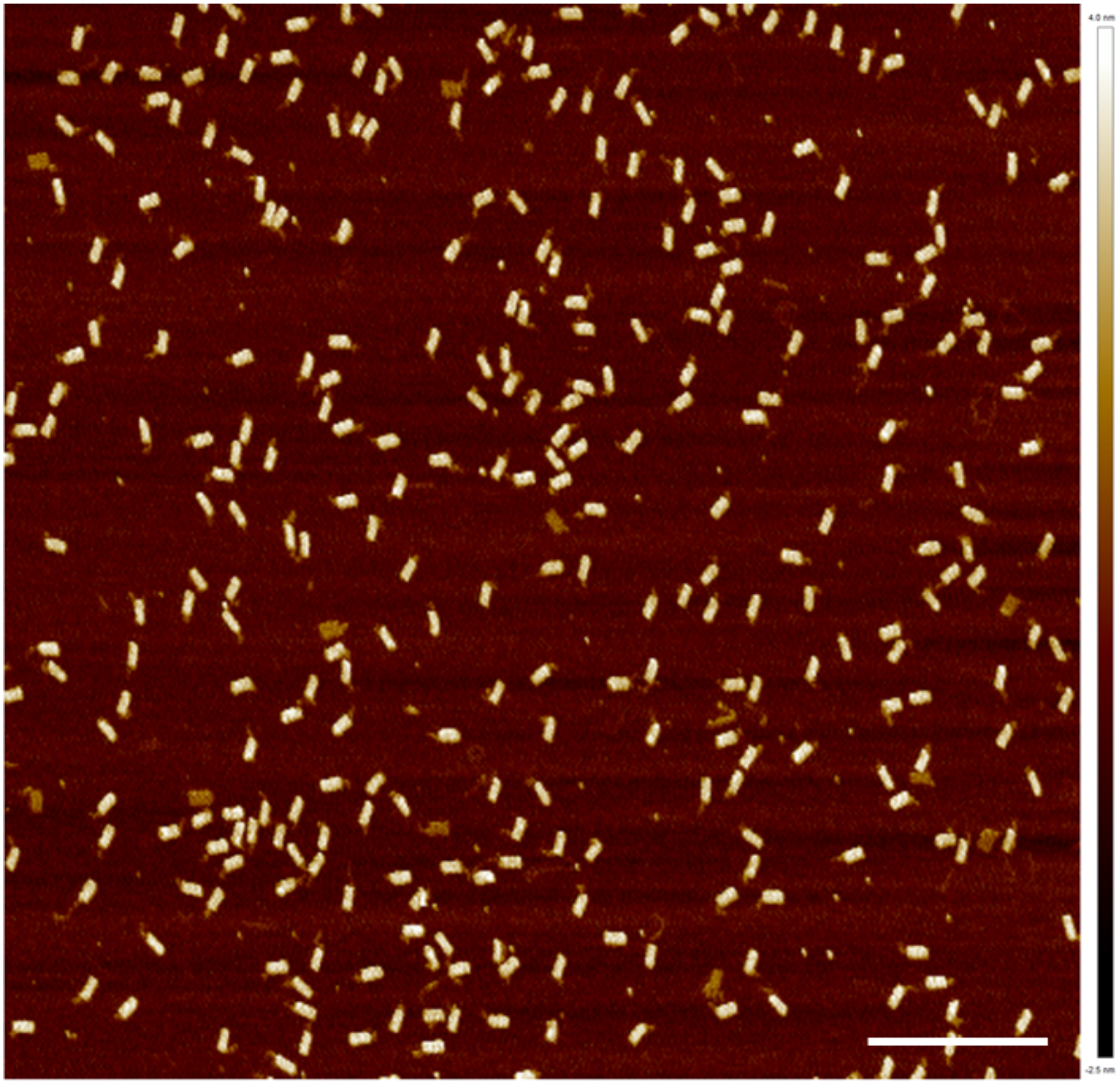


**Figure S4: Representative AFM image of the 18 helix structure (complex; GFP/mCherry template).** Sample was gel purified. Scale bar: 500 nm.

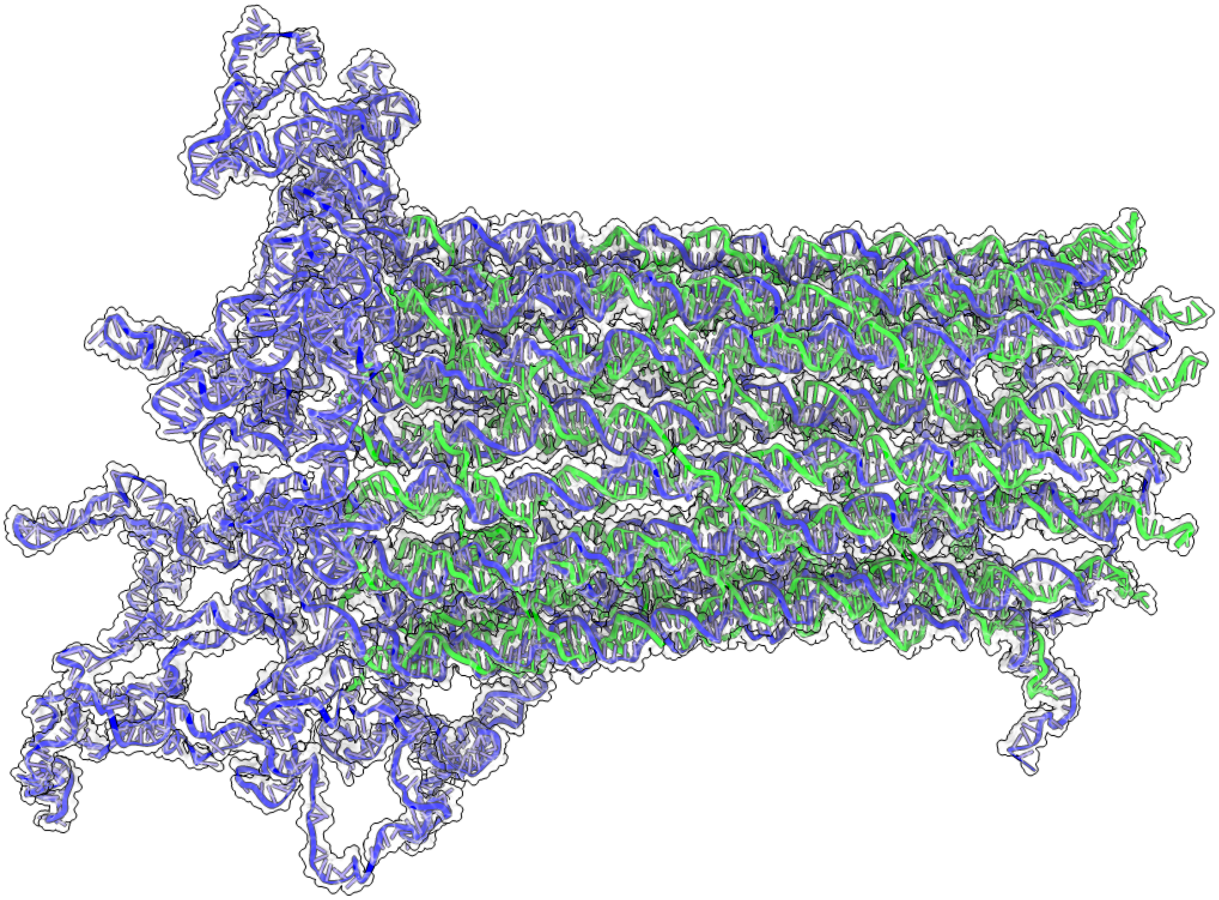

**Figure S5: oxDNA relaxation of the only top structure (GFP/mCherry template).**
Green color represents staple strands, blue color represents the template DNA. Simulations were run for 10^7 steps. ssDNA homology arms are not included to improve visualization of the structure.

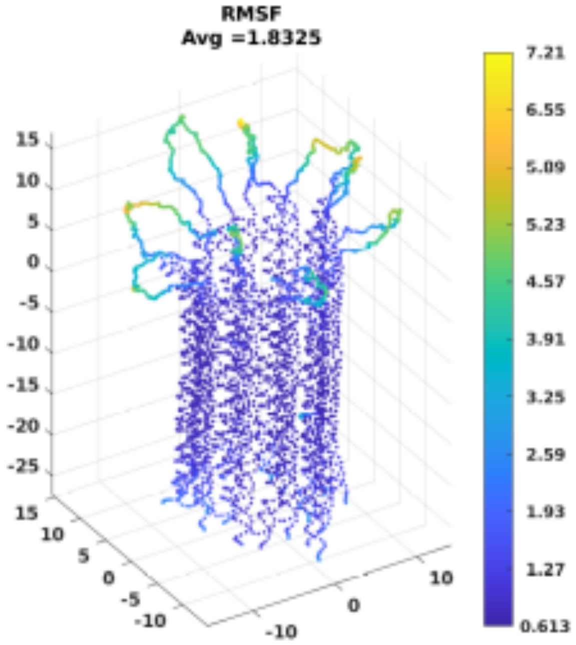


**Figure S6: Root-mean-square fluctuation analysis of oxDNA trajectories of the only top nanostructure (GFP/mCherry template).**

RMSF (in nm) of the only top nanostructure through 10^7 steps in oxDNA. Dark blue areas indicate smaller fluctuations than yellow areas. Processed in magicDNA.

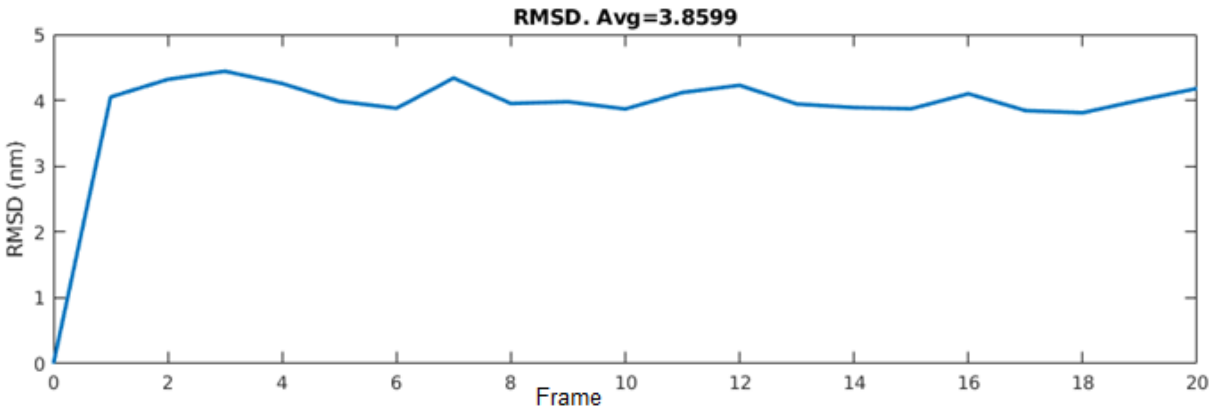


**Figure S7: Root-mean-square deviation analysis of oxDNA trajectories of the only topnanostructure (GFP/mCherry template).**

RMSD (in nm) of the 18 helix nanostructure through 10^7 steps in oxDNA. The graph indicates steady-state behavior as the RMSD reaches its plateau. Processed in magicDNA.

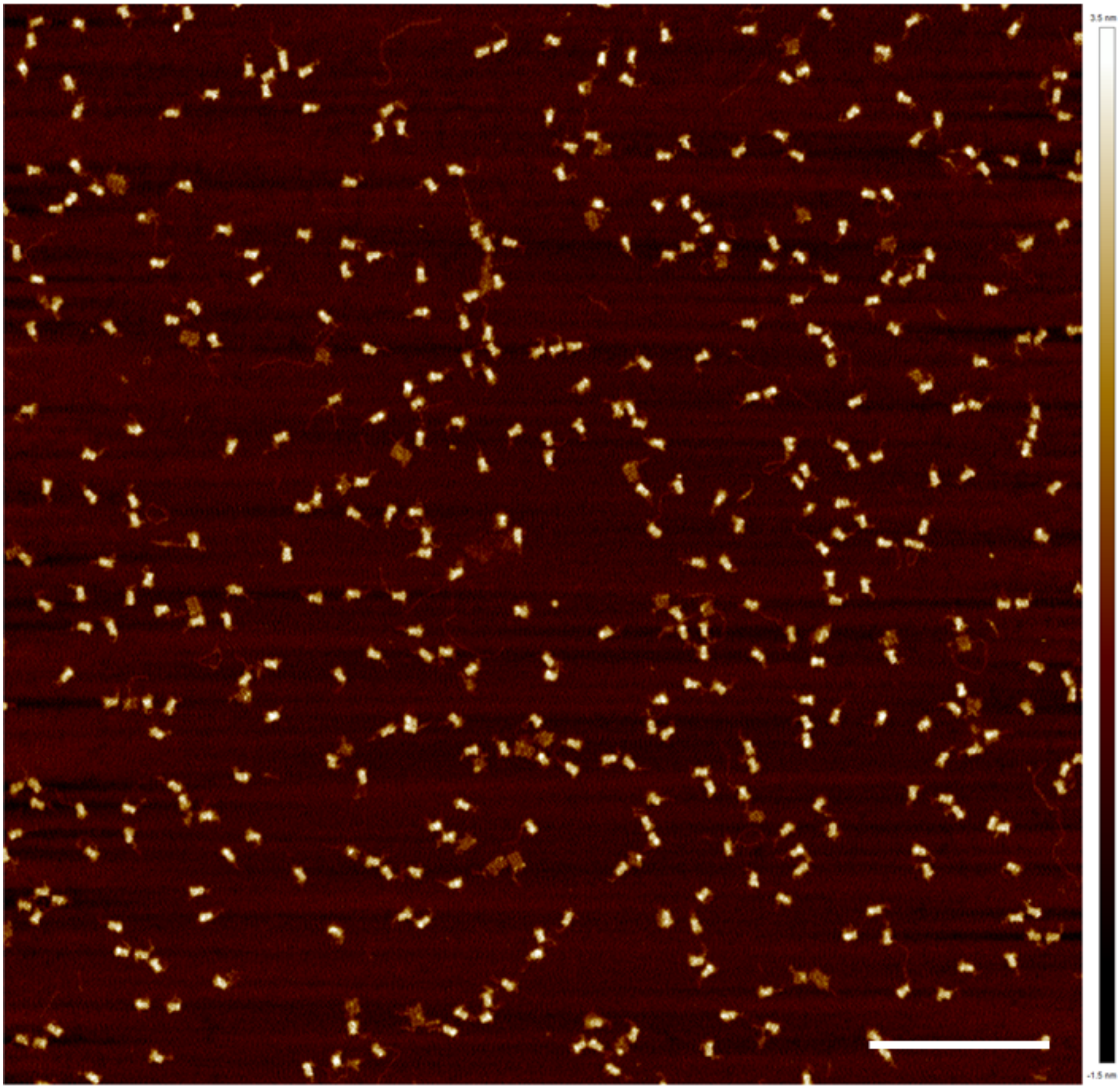

**Figure S8: Representative AFM image of the only top nanostructure (GFP/mCherry template).** Sample was gel purified. Scale bar: 500 nm.

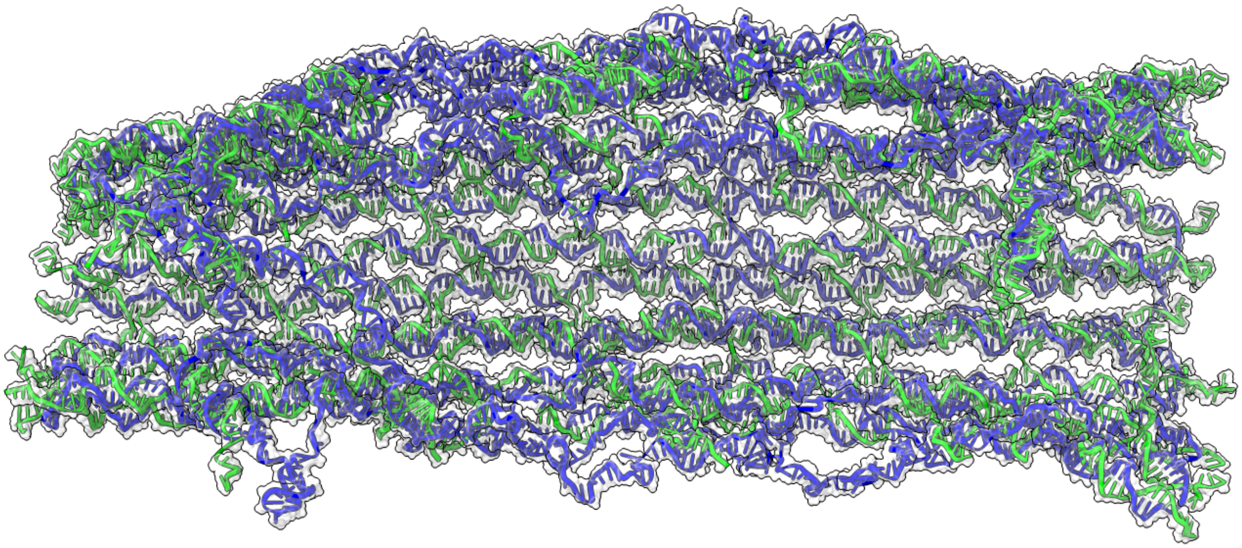

**Figure S9: oxDNA relaxation of the open nanostructure (GFP/mCherry template).**
Green color represents staple strands, blue color represents the template DNA. Simulations were run for 10^7 steps. ssDNA homology arms are not included to improve visualization of the structure.

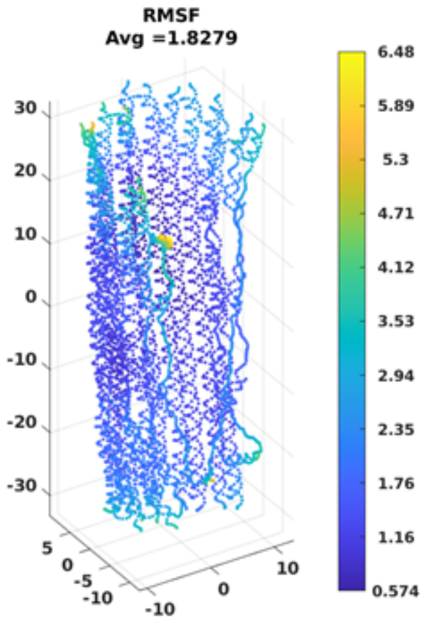


**Figure S10: Root-mean-square fluctuation analysis of oxDNA trajectories of the open nanostructure (GFP/mCherry template).**
RMSF (in nm) of the only top nanostructure through 10^7 steps in oxDNA. Dark blue areas indicate smaller fluctuations than yellow areas. Processed in magicDNA.

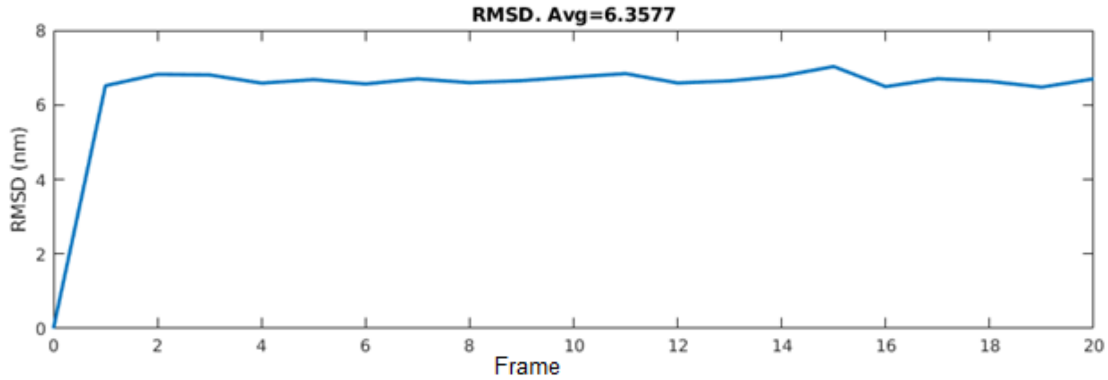


**Figure S11: Root-mean-square deviation analysis of oxDNA trajectories of the open nanostructure (GFP/mCherry template).**

RMSD (in nm) of the open nanostructure through 10^7 steps in oxDNA. The graph indicates steady-state behavior as the RMSD reaches its plateau. Processed in magicDNA.

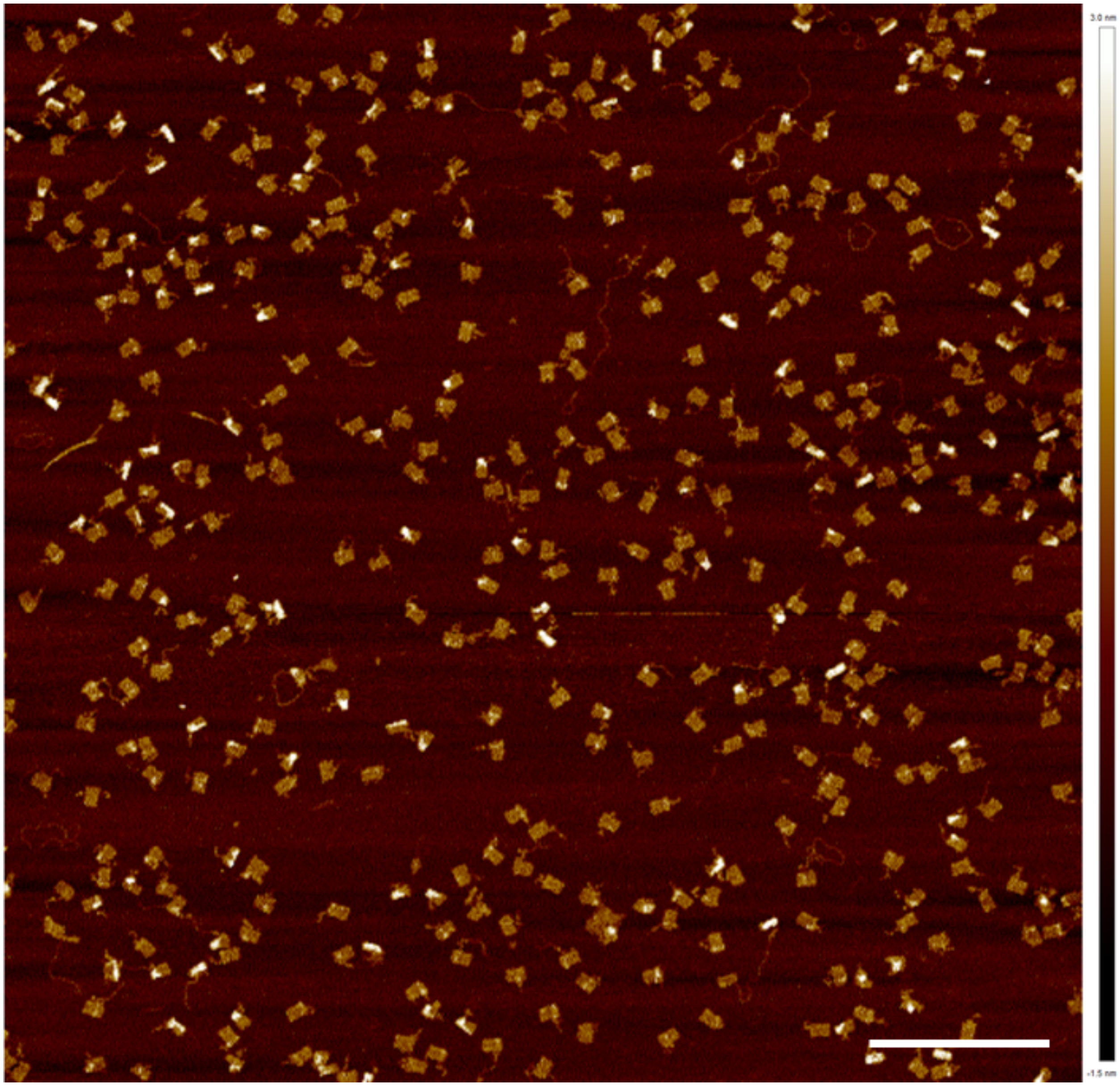


**Figure S12: Representative AFM image of the open nanostructure (GFP/mCherry template).** Sample was gel purified. Scale bar: 500 nm.

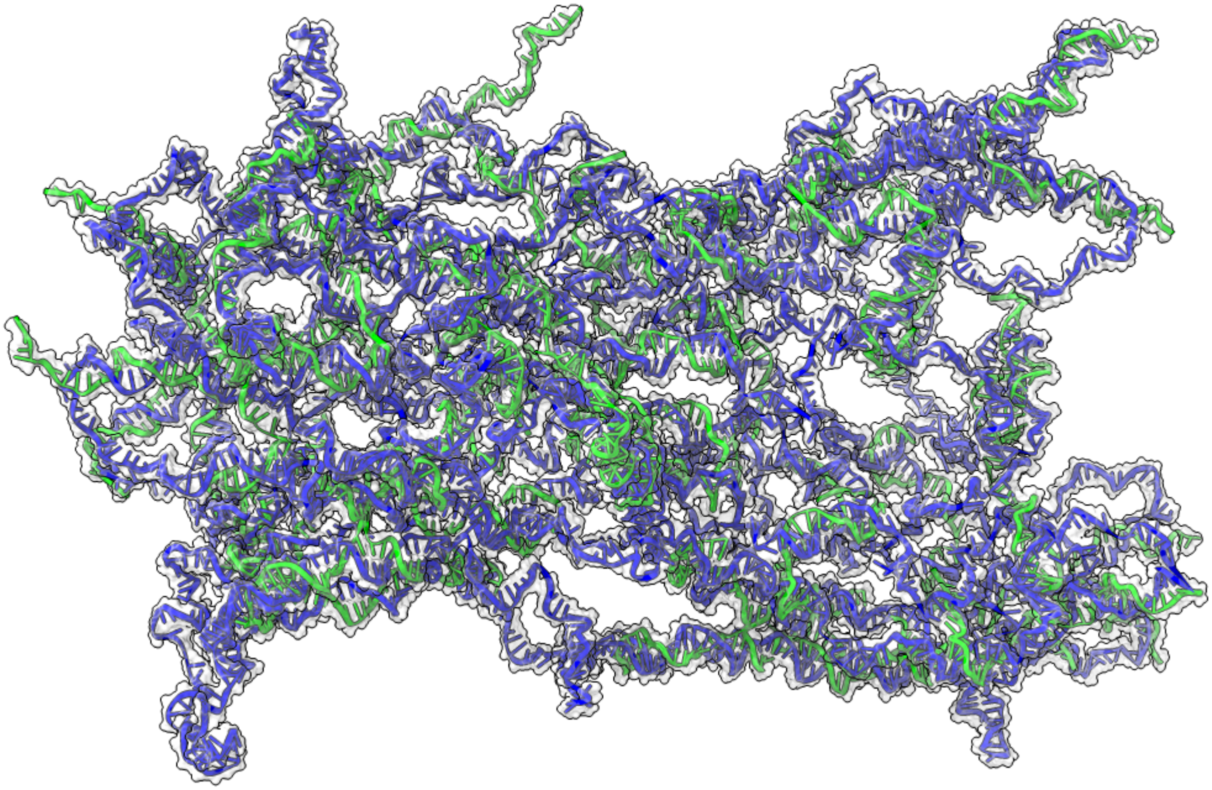

**Figure S13: oxDNA relaxation of the 50% staples nanostructure (GFP/mCherry template).** Green color represents staple strands, blue color represents the template DNA. Simulations were run for 10^7 steps. ssDNA homology arms are not included to improve visualization of the structure.

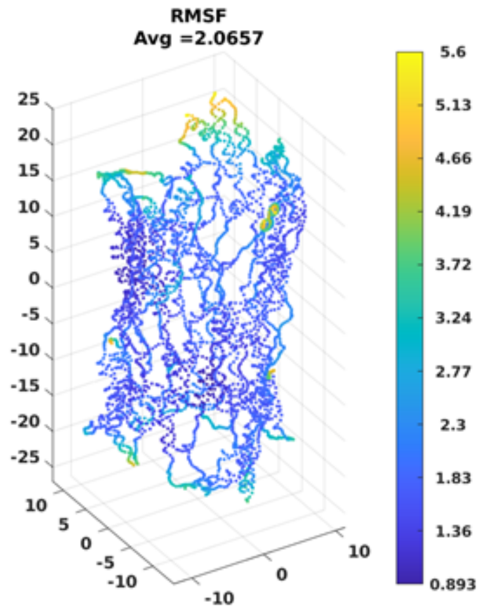


**Figure S14: Root-mean-square fluctuation analysis of oxDNA trajectories of the 50% staples nanostructure (GFP/mCherry template).**

RMSF (in nm) of the 50% nanostructure through 10^7 steps in oxDNA. Dark blue areas indicate smaller fluctuations than yellow areas. Processed in magicDNA.

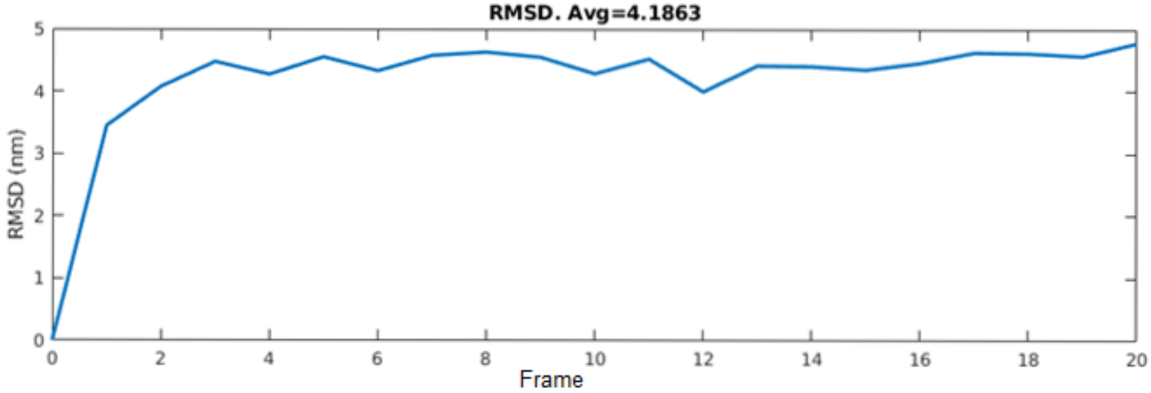


**Figure S15: Root-mean-square deviation analysis of oxDNA trajectories of the 50% staples nanostructure (GFP/mCherry template).**

RMSD (in nm) of the 50% nanostructure through 10^7 steps in oxDNA. The graph indicates steady-state behavior as the RMSD reaches its plateau. Processed in magicDNA.

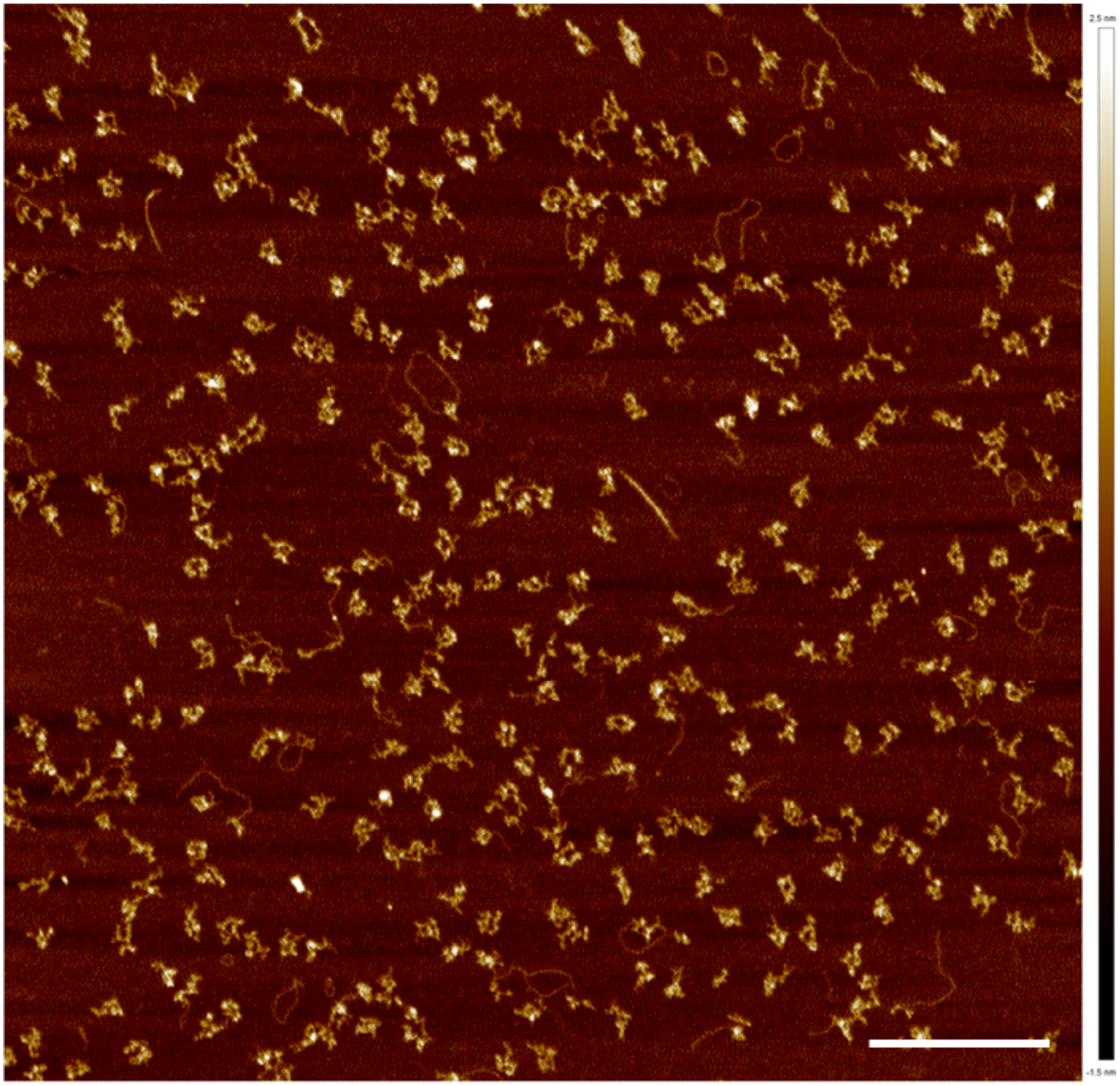

**Figure S16: Representative AFM image of the 50% staples nanostructure (GFP/mCherry template).** Sample was gel purified. Scale bar: 500 nm.

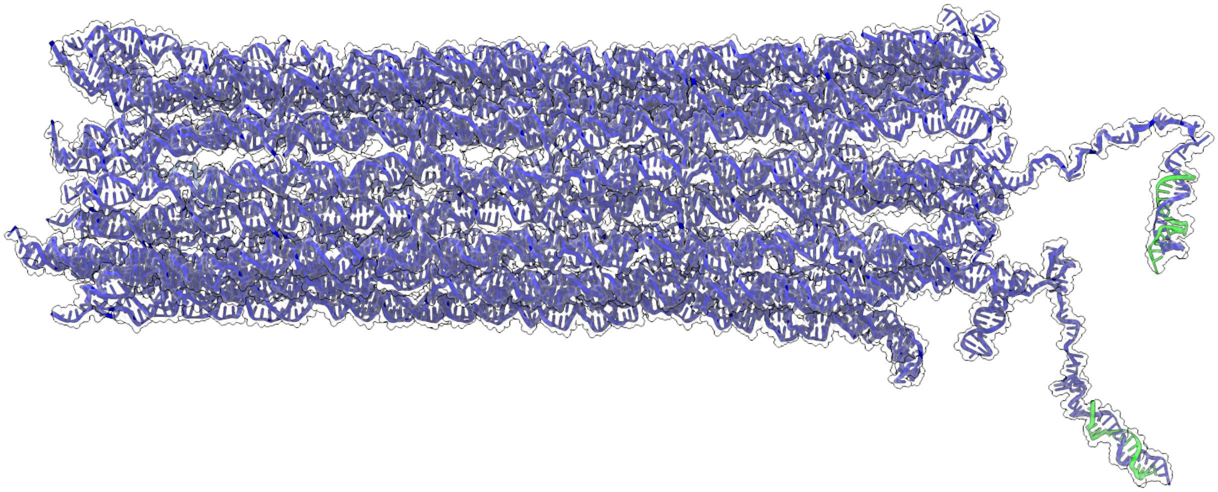

**Figure S17: oxDNA relaxation of the 18 helix nanostructure (complex), including ssDNA homology arms (mNeon template).**
Green color represents the shuttle sites, blue color represents the template DNA and staple strands used for structural formation. Simulations were run for 6.7 x 10^7 steps.

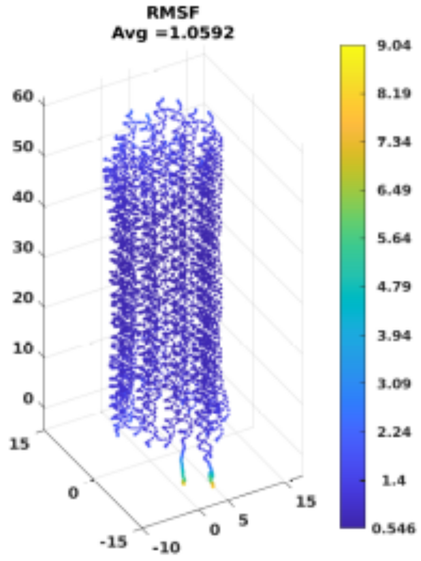


**Figure S18: Root-mean-square fluctuation analysis of oxDNA trajectories of the 18 helix nanostructure (complex), including ssDNA homology arms (mNeon template).**
RMSF (in nm) of the 18 helix nanostructure, including ssDNA homology arms (mNeon template) through 6.7 x 10^7 steps in oxDNA. Dark blue areas indicate smaller fluctuations than yellow areas. Processed in magicDNA.

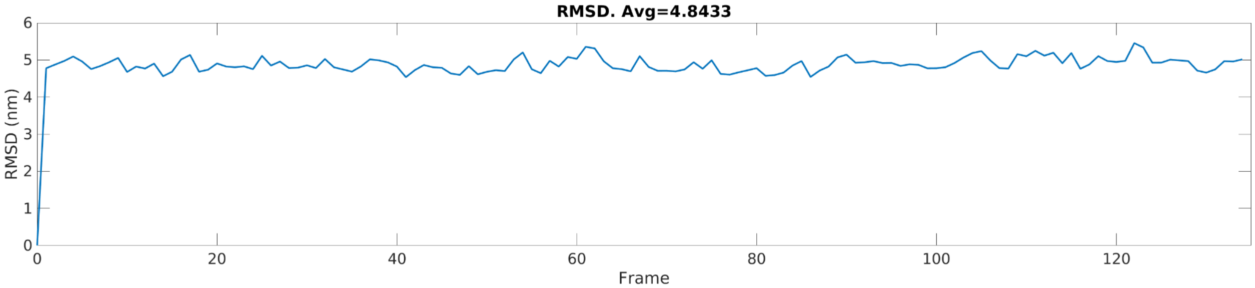


**Figure S19: Root-mean-square deviation analysis of oxDNA trajectories of the 18 helix nanostructure (complex), including ssDNA homology arms (mNeon template).**
RMSD (in nm) of the 18 helix nanostructure, including ssDNA homology arms (mNeon template) through 6.7 x 10^7 steps in oxDNA. The graph indicates steady-state behavior as the RMSD reaches its plateau. Processed in magicDNA.

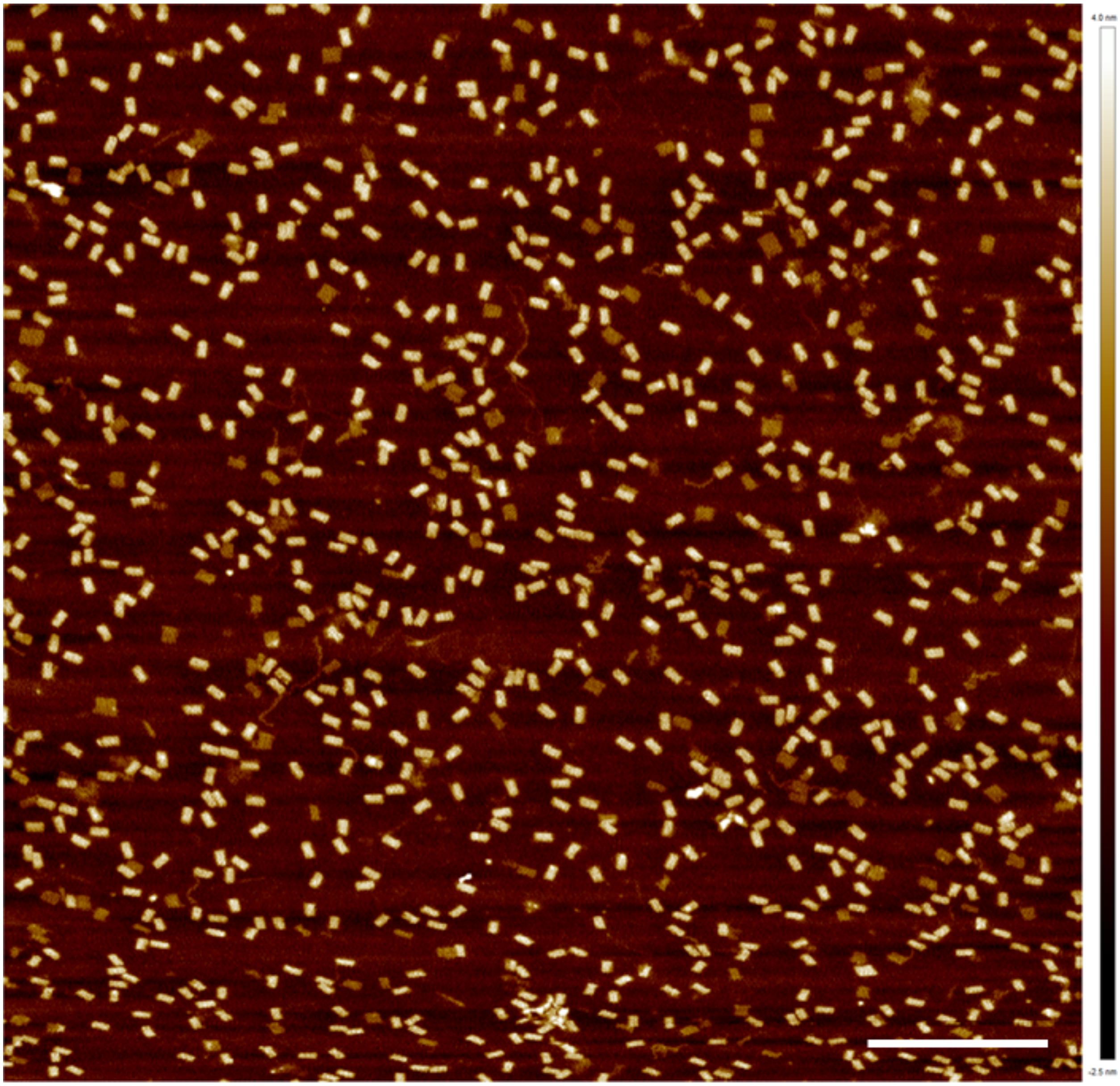

**Figure S20: Representative AFM image of the 18 helix nanostructure (complex) (mNeon template).** Sample was gel purified. Scale bar: 500 nm.

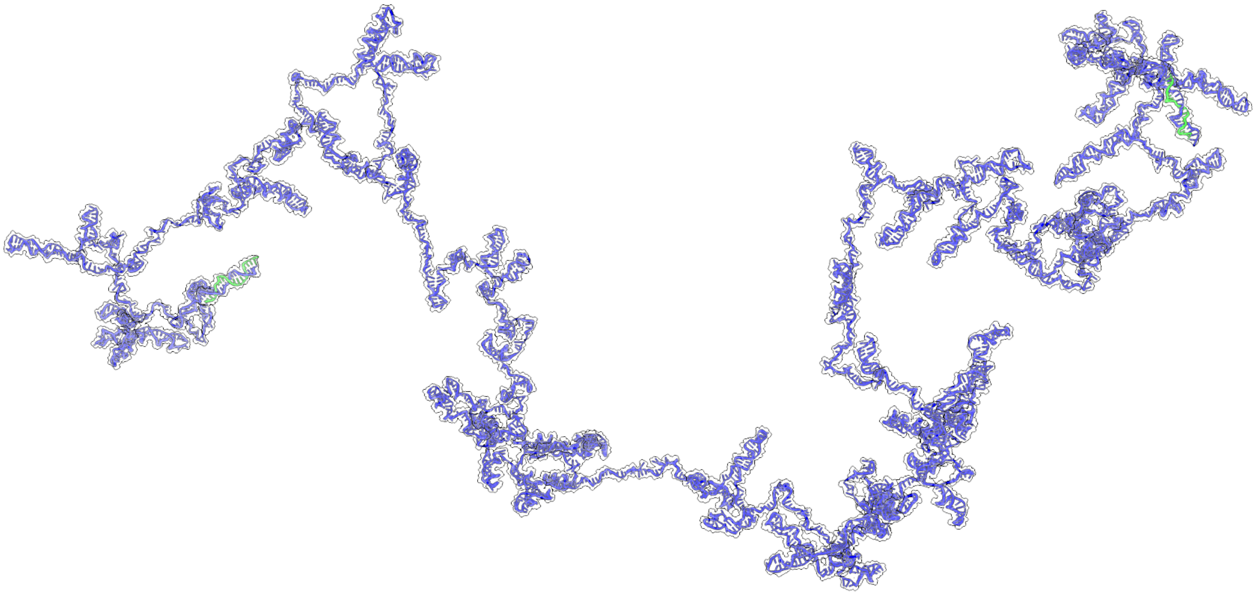

**Figure S21: oxDNA relaxation of the unstructured ssDNA (GFP/mCherry template).** Green color represents the shuttle sites, blue color represents the template DNA. Simulations were run for 1.1 x 10^8 steps.

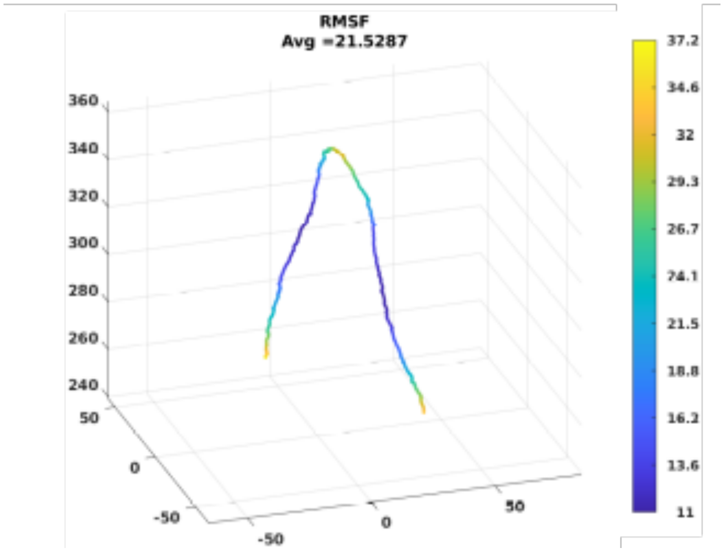


**Figure S22: Root-mean-square fluctuation analysis of oxDNA trajectories of the unstructured ssDNA (GFP/mCherry template).**

RMSF (in nm) of the unstructured ssDNA through 1.1 x 10^8 steps in oxDNA. Dark blue areas indicate smaller fluctuations than yellow areas. Processed in magicDNA. The color-coded visualization of the RMSF is based on an initial configuration and does not represent the simulation result.

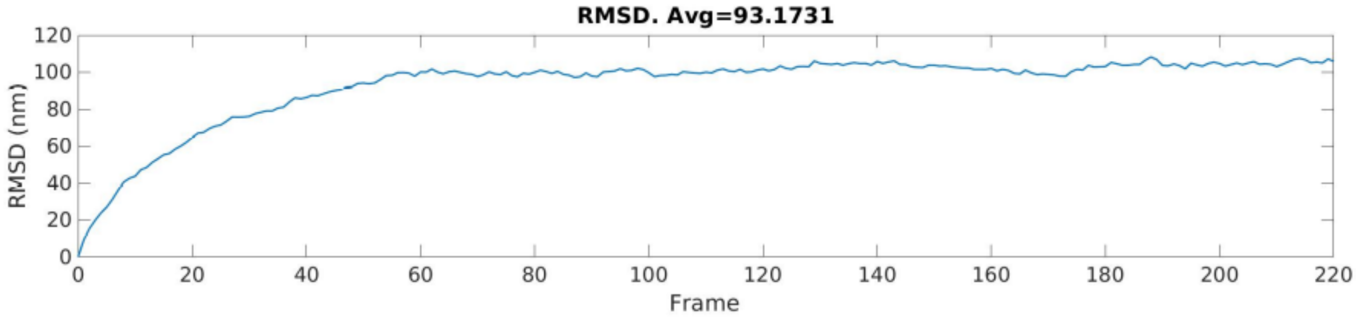


**Figure S23: Root-mean-square deviation analysis of oxDNA trajectories of the unstructured ssDNA (GFP/mCherry template).**
RMSD (in nm) of the unstructured ssDNA through 1.1 x 10^8 steps in oxDNA. The graph indicates steady-state behavior as the RMSD reaches its plateau. Processed in magicDNA.

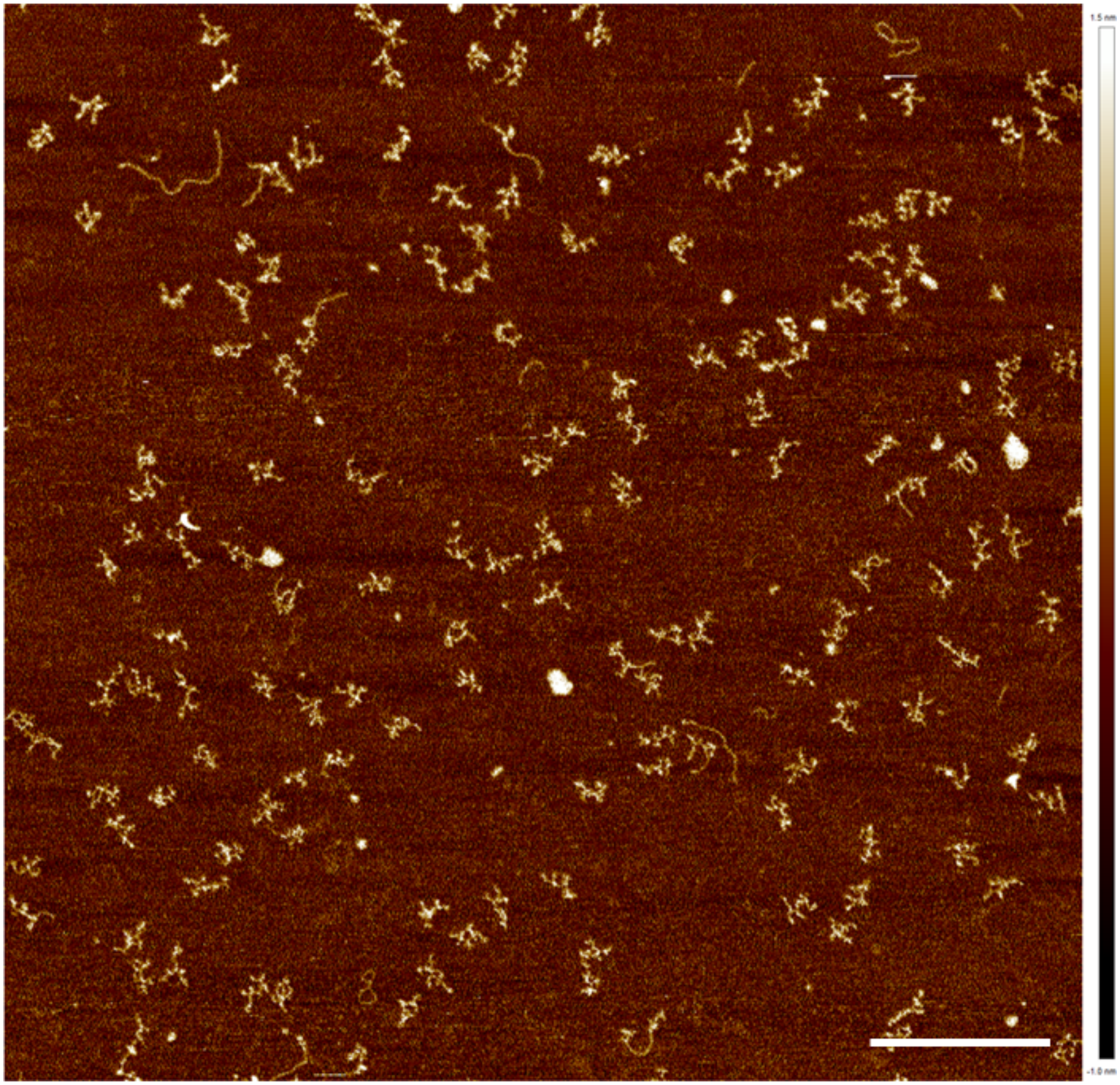

**Figure S24: Representative AFM image of the unstructured ssDNA (GFP/mCherry template).** Sample was gel purified. Scale bar: 500 nm.

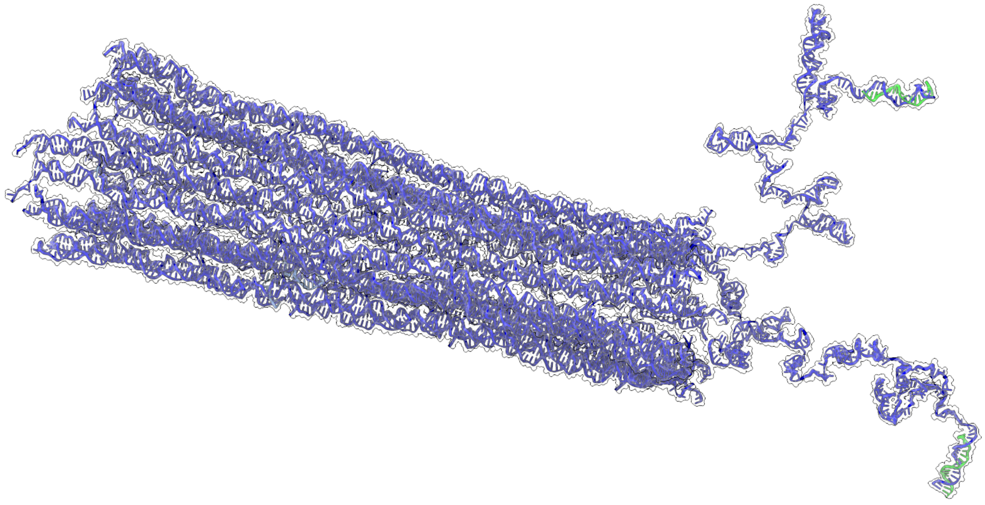

**Figure S25: oxDNA relaxation of the 18 helix nanostructure (complex), including ssDNA homology arms (GFP/mCherry template).**
Green color represents the shuttle sites, blue color represents the template DNA and staple strands used for structural formation. Simulations were run for 1.1 x 10^8 steps.

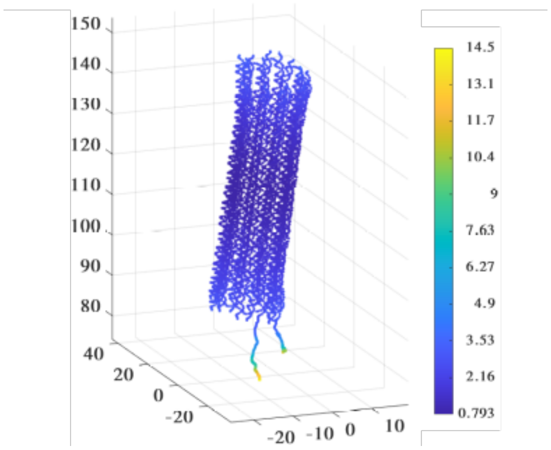


**Figure S26: Root-mean-square fluctuation analysis of oxDNA trajectories of the 18 helix nanostructure (complex), including ssDNA homology arms (GFP/mCherry template).** RMSF (in nm) of the 18 helix nanostructure, including ssDNA homology arms through 1.1 x 10^8 steps in oxDNA. Dark blue areas indicate smaller fluctuations than yellow areas. Processed in magicDNA.

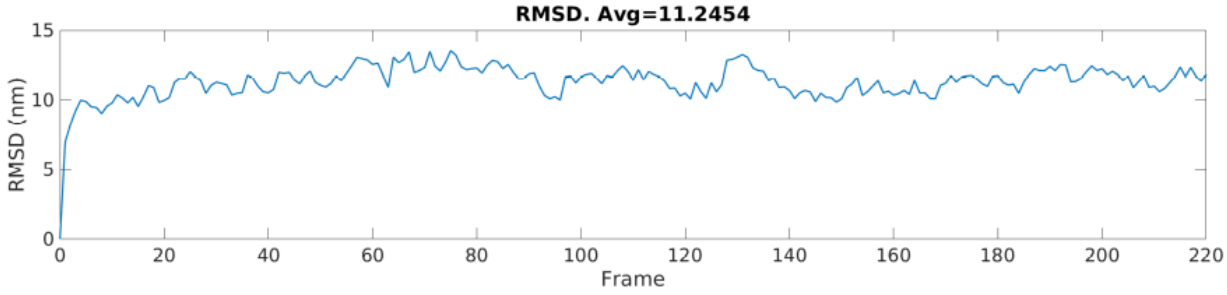


**Figure S27: Root-mean-square deviation analysis of oxDNA trajectories of the 18 helix nanostructure (complex), including ssDNA homology arms (GFP/mCherry template).**
RMSD (in nm) of the 18 helix nanostructure, including ssDNA homology arms through 10^7 steps in oxDNA. The graph indicates steady-state behavior as the RMSD reaches its plateau. Processed in magicDNA.

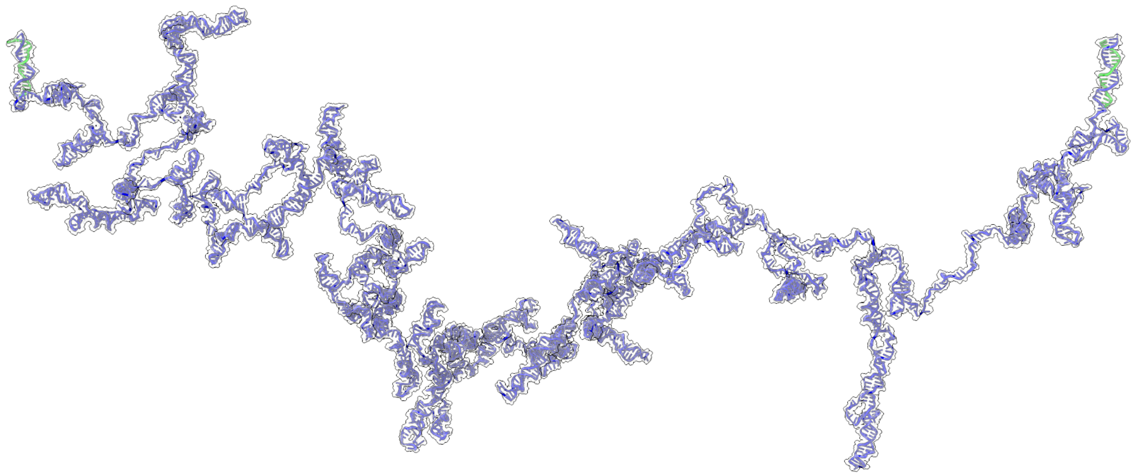

**Figure S28: oxDNA relaxation of the unstructured ssDNA (mNeon template).**
Green color represents the shuttle sites, blue color represents the template DNA. Simulations were run for 2 x 10^8 steps. The color-coded visualization of the RMSF is based on an initial configuration and does not represent the simulation result.

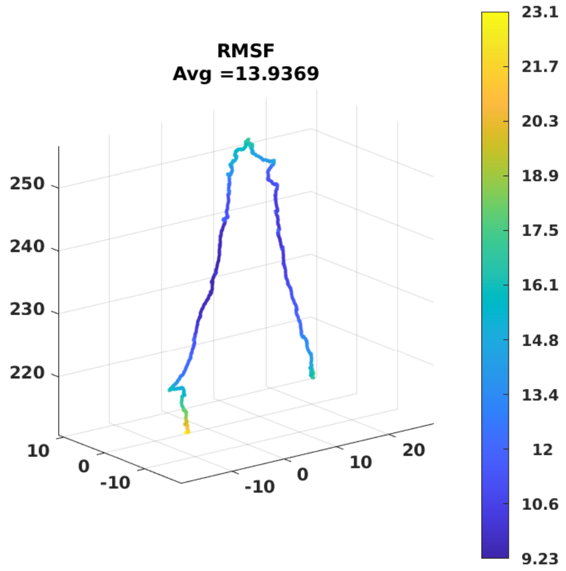


**Figure S29: Root-mean-square fluctuation analysis of oxDNA trajectories of the unstructured ssDNA (mNeon template).**

RMSF (in nm) of the unstructured ssDNA through 2 x 10^8 steps in oxDNA. Dark blue areas indicate smaller fluctuations than yellow areas. Processed in magicDNA. The color-coded visualization of the RMSF is based on an initial configuration and does not represent the simulation result.

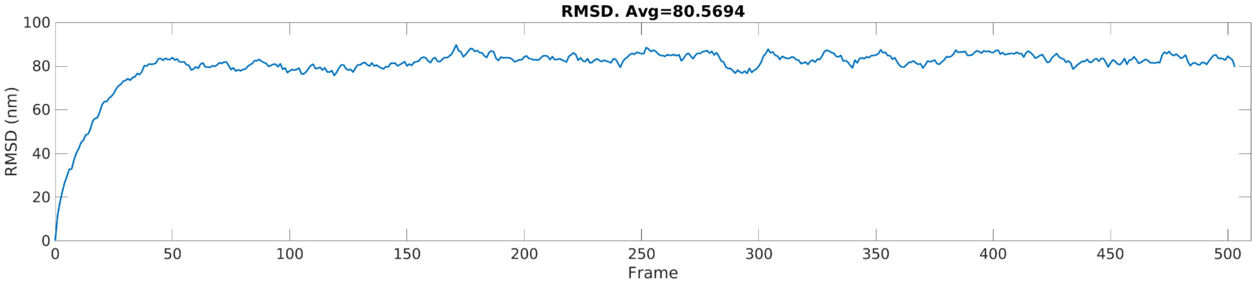


**Figure S30: Root-mean-square deviation analysis of oxDNA trajectories of the unstructured ssDNA (mNeon template).**

RMSD (in nm) of the unstructured ssDNA through 2 x 10^8 steps in oxDNA. The graph indicates steady-state behavior as the RMSD reaches its plateau. Processed in magicDNA.

**Shuttle-site distance measurements**

We used a custom python code (*50*) to determine the average distance between the two shuttle-sites for the 18 helix full structure and the unstructured ssDNA.

We ran simulations between 130 and 503 frames and used data after the RMSD reached its plateau to prevent the initial configuration from affecting our results.

Figures S17, 21, 25, 28 show representative configurations of all structures, with highlighted shuttle oligos. Supplementary movies 5 - 8 show the trajectories of these simulations.

We measured the distance between the last base of the shuttle sequence over time.

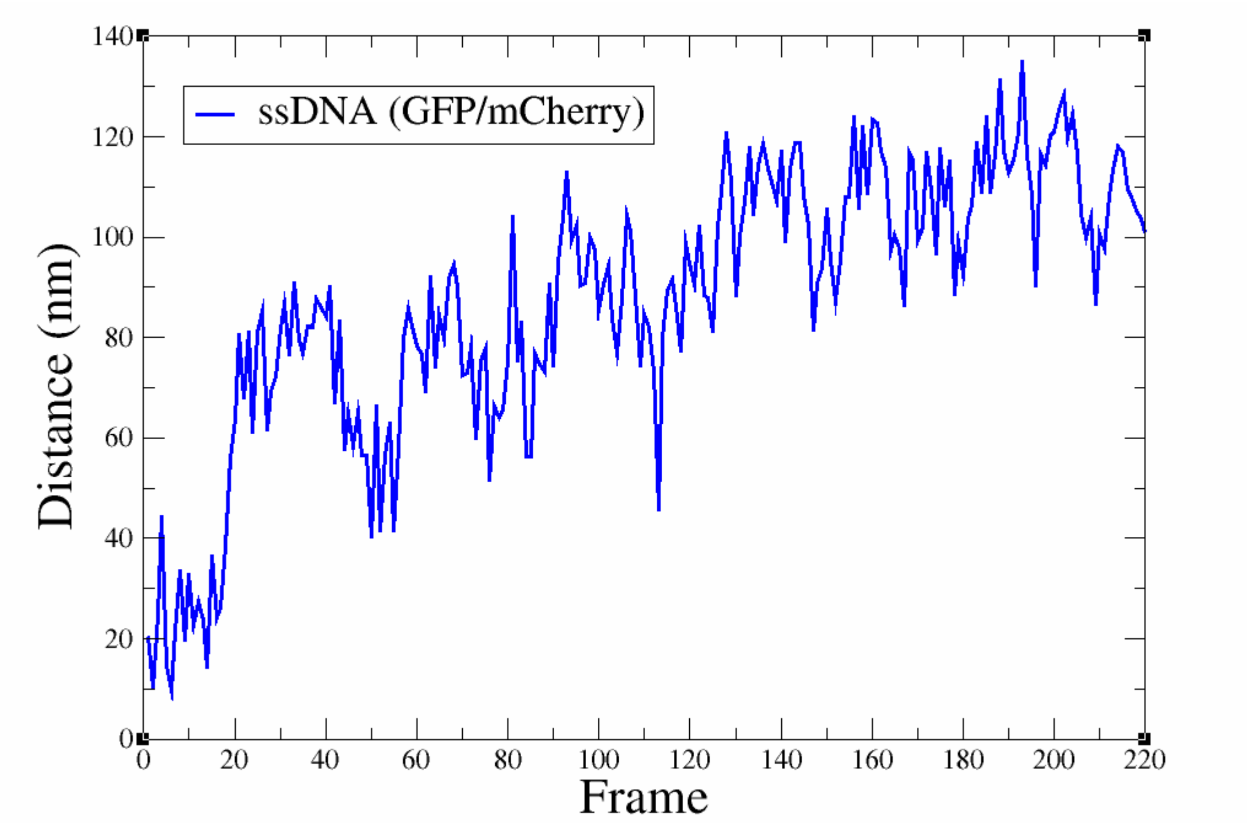


**Figure S31: Distances between the shuttle sites on the homology arms of the unstructured ssDNA (GFP/mCherry template) by oxDNA.** Simulations were run for a total of 220 frames, the average distance between both shuttle sites (after reaching a RMSD plateau) is 108.98 ± 11.22 nm.

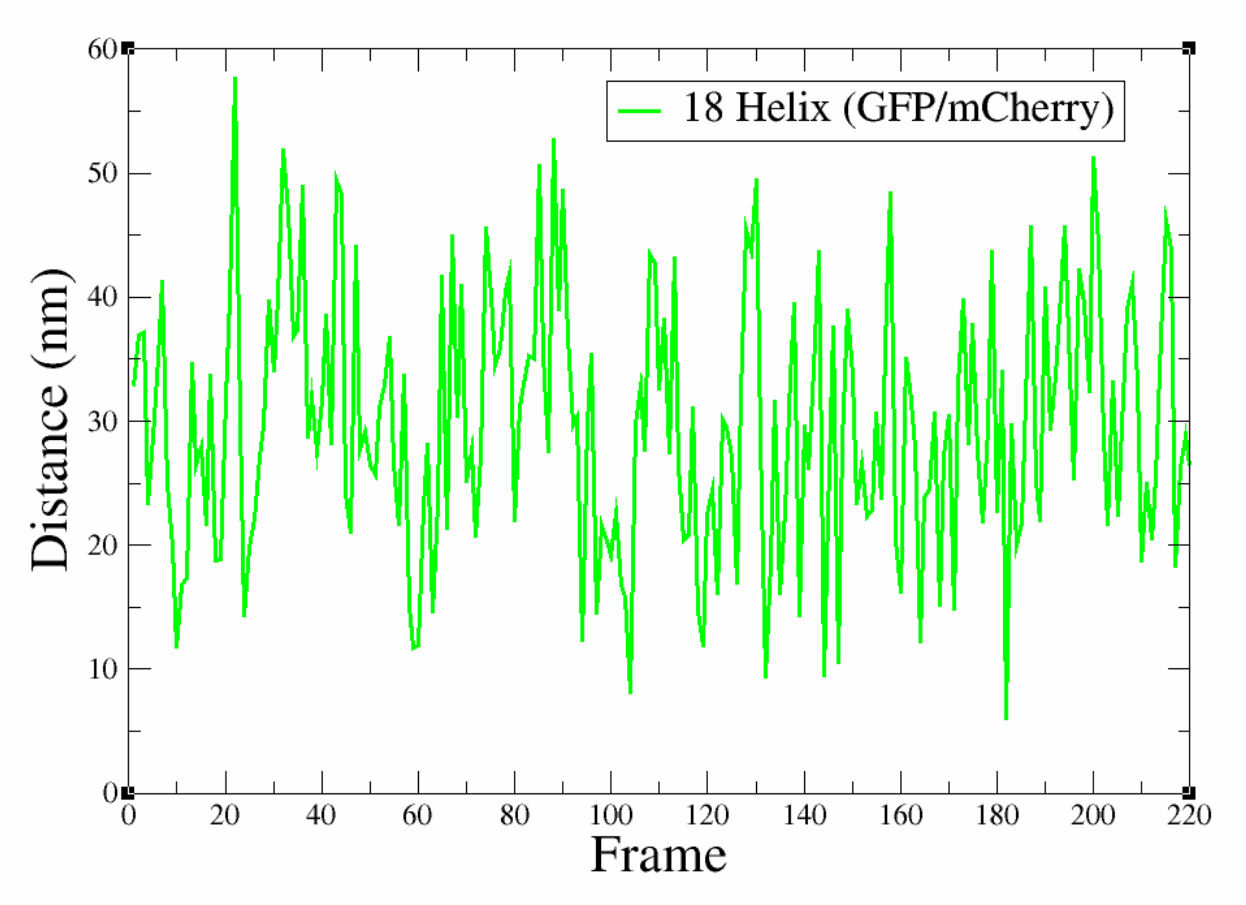


**Figure S32: Distances between the shuttle sites on the homology arms of the 18 helix nanostructure (complex; GFP/mCherry template) by oxDNA.** Simulations were run for a total of 220 frames, the average distance between both shuttle sites (after reaching a RMSD plateau) is 29.33 ± 9.9 nm.

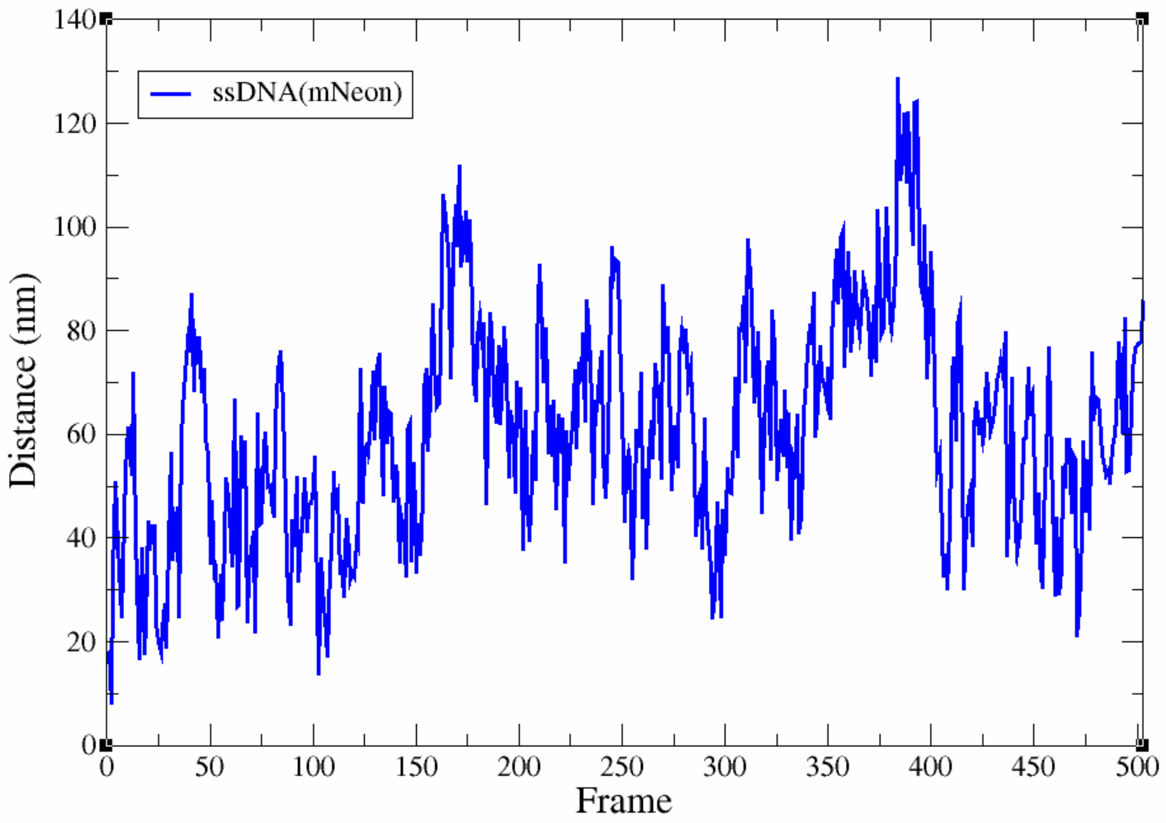


**Figure S33: Distances between the shuttle sites on the homology arms of the unstructured ssDNA (mNeon template) by oxDNA.** Simulations were run for a total of 500 frames, the average distance between both shuttle sites (after reaching a RMSD plateau) is 65.67 ± 19.71 nm.

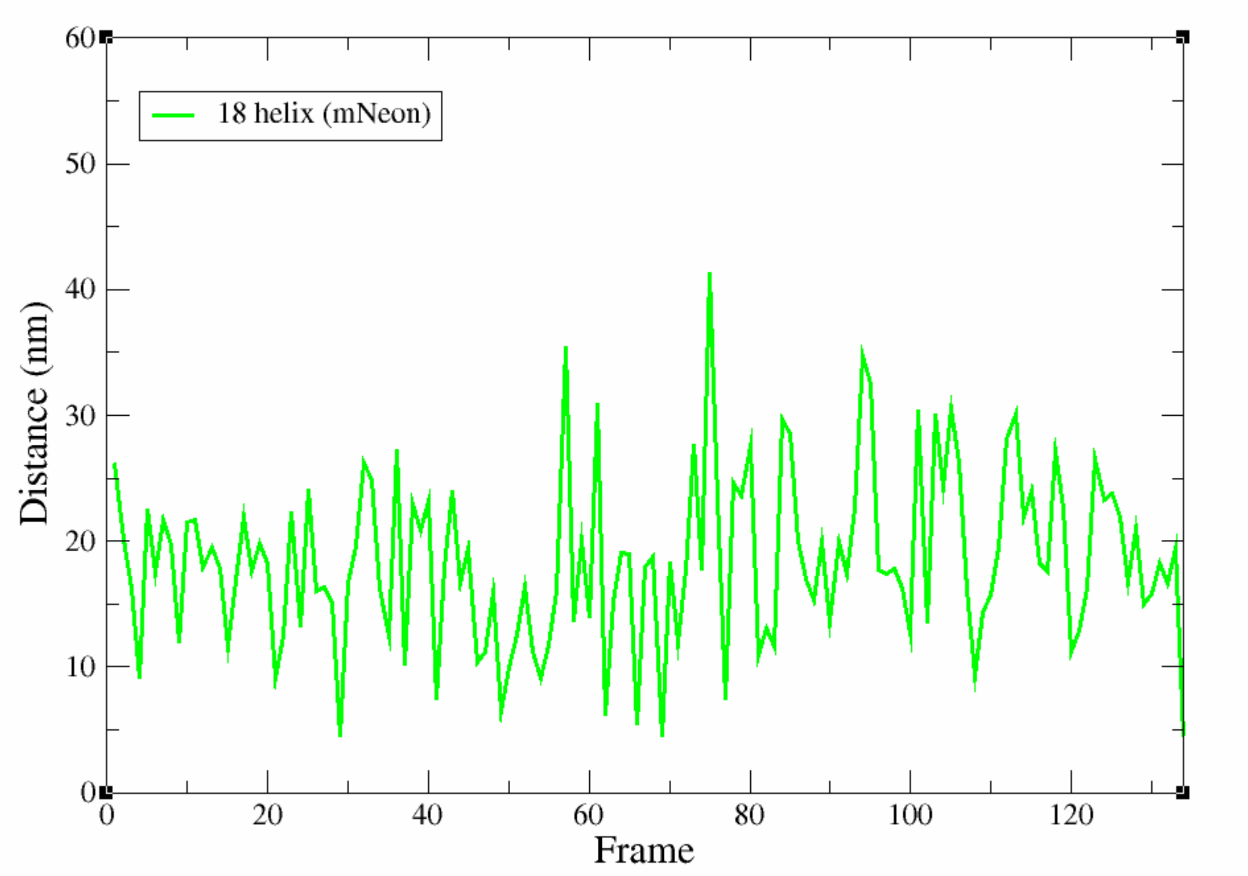


**Figure S34: Distances between the shuttle sites on the homology arms of the 18 helix nanostructure (complex; mNeon template) by oxDNA.** Simulations were run for a total of 130 frames, the average distance between both shuttle sites (after reaching a RMSD plateau) is 19.7 ± 7.2 nm.

**Table S3: Average distance between shuttle sites for 18 helix nanostructures and unstructured ssDNA on both HDR templates.**

| Structure Name | Average Shuttle Distance [nm] | Standard Deviation [nm] | Frames Used for Calculations |
|---|---|---|---|
| ssDNA (GFP/mCherry) | 108.98 | 11.22 | 130-220 |
| 18 Helix (GFP/mCherry) | 29.33 | 9.9 | 130-220 |
| ssDNA (mNeon) | 65.67 | 19.71 | 220-503 |
| 18 Helix (mNeon) | 19.7 | 7.2 | 65-134 |

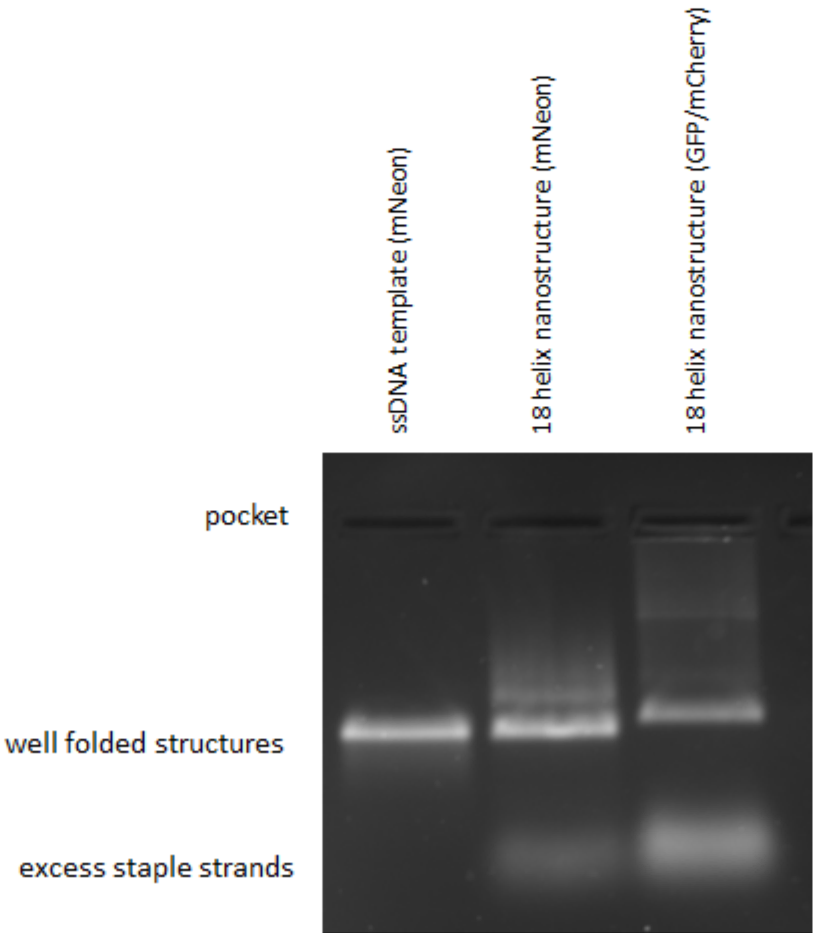


**Figure S35: Agarose gel electrophoresis of 18 helix nanostructures on both templates.** From left to right: ssDNA template (mNeon), 18 helix nanostructure on the mNeon template, 18 helix nanostructure on the GFP/mCherry template. The lower electrophoretic mobility of the 18 helix nanostructure on the GFP/mCherry template can most likely be attributed to the longer ssDNA homology arms.

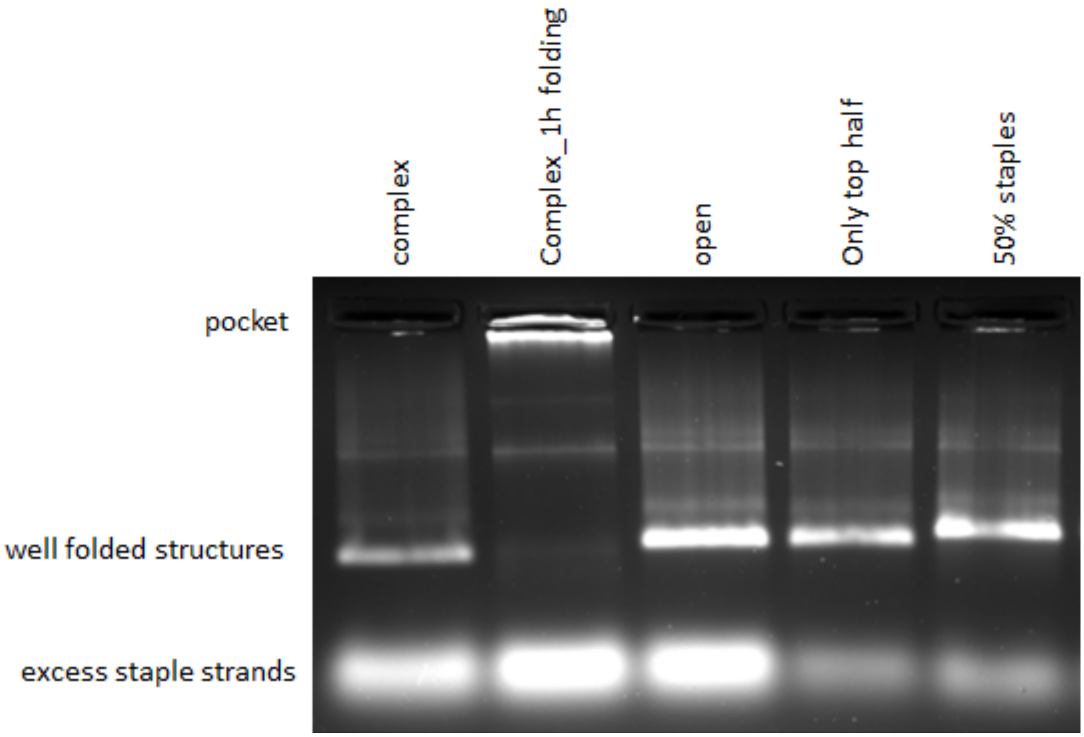


**Figure S36: Agarose gel electrophoresis of folded structures on the GFP/mCherry template.** From left to right: 18 helix nanostructure, 18 helix nanostructure, folded with a faster thermal annealing protocol, the open nanostructure, the only top nanostructure and the 50% staples nanostructure. The 18 helix nanostructure folded with a faster thermal annealing protocol does not fold properly, as indicated by the strong signal in the gel pocket and is not further discussed in the main text.

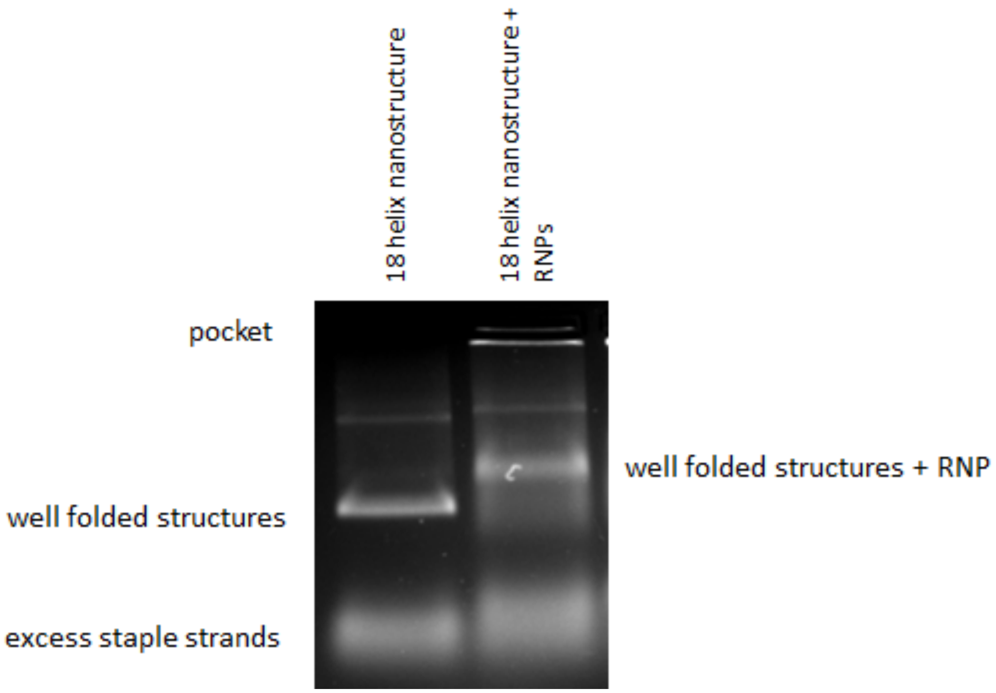


**Figure S37: Agarose gel electrophoresis of the 18 helix structure (GFP/mCherry) with RNPs.** Left band: The 18 helix nanostructure without RNPs bound. Right band: The 18 helix nanostructure was pre-incubated for 10 minutes at room temperature with RNPs prior to gel electrophoresis.

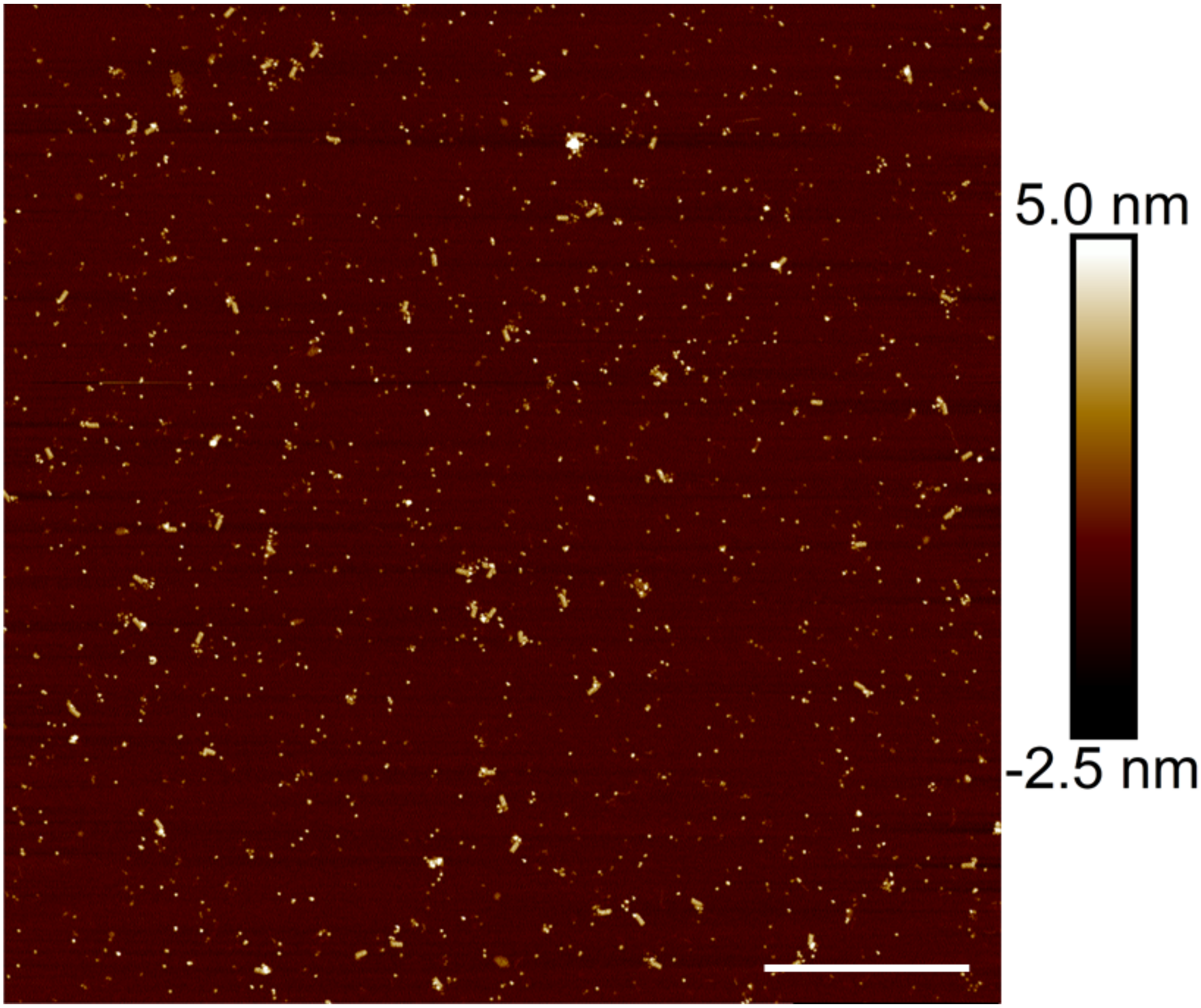


**Figure S38: Representative AFM image of the 18 helix nanostructure with RNPs (GFP/mCherry template).** Sample was gel purified. Scale bar: 1 µm.
